# Clinical recovery of *Macaca fascicularis* infected with *Plasmodium knowlesi*

**DOI:** 10.1101/2021.06.28.448877

**Authors:** Mariko S. Peterson, Chester J. Joyner, Jessica A. Brady, Jennifer S. Wood, Monica Cabrera-Mora, Celia L. Saney, Luis L. Fonseca, Wayne T. Cheng, Jianlin Jang, Stacey A. Lapp, Stephanie R. Soderberg, Mustafa V. Nural, Jay C. Humphrey, Allison Hankus, Deepa Machiah, Ebru Karpuzoglu, Jeremy D. DeBarry, MaHPIC-Consortium, Rabindra Tirouvanziam, Jessica C. Kissinger, Alberto Moreno, Sanjeev Gumber, Eberhard O. Voit, Juan B. Gutiérrez, Regina Joice Cordy, Mary R. Galinski

**Affiliations:** Yerkes National Primate Research Center, Emory University, Atlanta, GA, USA; Emory Vaccine Center, Emory University, Atlanta, GA, USA; Center for Vaccines and Immunology, Department of Infectious Diseases, University of Georgia, Athens GA, USA; School of Chemical, Materials and Biomedical Engineering, University of Georgia, Athens, GA, USA; Division of Animal Resources, Yerkes National Primate Research Center, Emory University, Atlanta, GA, USA; The Wallace H. Coulter Department of Biomedical Engineering, Georgia Institute of Technology and Emory University, Atlanta, GA, USA; Institute of Bioinformatics, University of Georgia, Athens, GA, USA; Division of Pathology, Yerkes National Primate Research Center, Atlanta, GA, USA; Department of Pediatrics, Emory University School of Medicine, Atlanta, GA, USA; Department of Genetics, University of Georgia, Athens, GA, USA; Center for Tropical and Emerging Global Diseases, University of Georgia, Athens, GA, USA; Division of Infectious Diseases, Department of Medicine, Emory University School of Medicine, Atlanta, GA; Department of Pathology and Laboratory Medicine, Emory School of Medicine, Atlanta, GA; Department of Mathematics, University of Texas at San Antonio, San Antonio, TX, USA

**Keywords:** Malaria, nonhuman primate models, infectious diseases, resilience, telemetry, fever, haematology, anaemia, thrombocytopaenia, erythropoietin, bone marrow, histopathology

## Abstract

**Background:** Kra monkeys (*Macaca fascicularis*), a natural host of *Plasmodium knowlesi,* control parasitaemia caused by this parasite species and escape death without treatment. Knowledge of the disease progression and resilience in kra monkeys will aid the effective use of this species to study mechanisms of resilience to malaria. This longitudinal study aimed to define clinical, physiological and pathological changes in kra monkeys infected with *P. knowlesi,* which could explain their resilient phenotype.

**Methods:** Kra monkeys (*n* = 15, male, young adults) were infected intravenously with cryopreserved *P. knowlesi* sporozoites and the resulting parasitaemias were monitored daily. Complete blood counts, reticulocyte counts, blood chemistry and physiological telemetry data (*n* = 7) were acquired as described prior to infection to establish baseline values and then daily after inoculation for up to 50 days. Bone marrow aspirates, plasma samples, and 22 tissue samples were collected at specific time points to evaluate longitudinal clinical, physiological and pathological effects of *P. knowlesi* infections.

**Results:** As expected, the kra monkeys controlled parasitaemia and remained with low-level, persistent parasitaemias without antimalarial intervention. Unexpectedly, early in the infection, fevers developed, which ultimately returned to baseline, as well as mild to moderate thrombocytopaenia, and moderate to severe anaemia. Mathematical modeling and the reticulocyte production index indicated that the anaemia was largely due to the removal of uninfected erythrocytes and not impaired production of erythrocytes. Mild tissue damage was observed, and tissue parasite load was associated with tissue damage even though parasite accumulation in the tissues was generally low.

**Conclusions:** Kra monkeys experimentally infected with *P. knowlesi* sporozoites presented with multiple clinical signs of malaria that varied in severity among individuals. Overall, the animals shared common mechanisms of resilience characterized by controlling parasitaemia 3-5 days after patency, and controlling fever, coupled with physiological and bone marrow responses to compensate for anaemia. Together, these responses likely minimized tissue damage while supporting the establishment of chronic infections, which may be important for transmission in natural endemic settings. These results provide new foundational insights into malaria pathogenesis and resilience in kra monkeys, which may improve understanding of human infections.

## BACKGROUND

*Plasmodium knowlesi* malaria cases have been identified in 10 of the 11 countries that comprise Southeast Asia; the one exception being Timor-Leste [reviewed in 1, 2, 3]. Since its recognition in 2004 in Malaysia as an emergent zoonotic parasite species [4], *P. knowlesi* has in fact risen as the major cause of human clinical cases of malaria in Malaysia and threatens to upend malaria elimination in this country [1, 2, 4–8]; over 4,000 Malaysian cases of *P. knowlesi* malaria were reported in 2018 [9]. *Plasmodium knowlesi* transmission to humans has been attributed to spillover events from peri-domestic infected macaques in village settings, or from infected wild macaques in the setting of jungle trekking, foraging, farming, and logging activities [1, 5, 10–13]. At least three macaque species are endemic to these geographical regions and can serve as reservoirs for *P. knowlesi*: *Macaca fascicularis* (the kra monkey, or long-tailed macaque), *Macaca nemestrina* (pig-tailed macaque) and *Macaca arctoides* (stump-tailed macaque) [14–17].

*Plasmodium knowlesi* infections cause symptoms that are generally associated with malaria caused by other *Plasmodium* species (*Plasmodium falciparum*, *Plasmodium vivax*, *Plasmodium malariae*, and *Plasmodium ovale,* and the closely related zoonotic species *Plasmodium cynomolgi*) [18–21], including cyclical episodes of fever and chills with rigor. Clinical complications have the potential to become exacerbated quickly given this species’ unique 24-hour replication cycle within the erythrocyte host cell [1, 5, 22–24]. Asymptomatic *P. knowlesi* infections are less studied, but they have been shown to be common, including a high percentage in children [12]. Clinically, thrombocytopaenia has been reported as a universal problem with *P. knowlesi* infections (<150,000 platelets/μl), with about 1/3 of adult patients showing severe thrombocytopaenia (<50,000 platelets/μl) [reviewed in 1]. Anaemia is not a predominant problem with *P. knowlesi* malaria in adults, though moderate anaemia (defined as less than 10 g/dL) and recovery from this disease state have been observed, and some cases of severe anaemia (less than 7 g/dL) have been documented [24, 25] and reviewed in [1]. In contrast, anaemia has been identified as a predominant feature in children with *P. knowlesi* malaria [26], despite children having lower parasitaemias (and no fatalities reported) [27]. While hyperparasitaemia is associated with severe disease, it should be noted that *P. knowlesi* can cause clinical manifestations at lower parasitaemias than other species (e.g., baseline median values on hospital admission in Kapit of 1,387 *P. knowlesi* parasites/μl, compared to 4,258 *P. vivax* parasites/μl and 26,781 *P. falciparum* parasites/μl) [24]. Also, Barber and others showed in a prospective study that in Saba, Malaysia, *P. knowlesi* was more likely than *P. falciparum* to cause severe disease [25].

In severe cases of *P. knowlesi* malaria, clinical signs and symptoms in adults have included abdominal pain, shortness of breath, productive cough, and respiratory distress [1, 6, 23, 24]. Severe cases have included jaundice, acute kidney injury, metabolic acidosis, acute lung injury, and shock, among others [reviewed in 28, 29]. High parasitaemia (>20,000 *P. knowlesi* parasites/μl) along with jaundice and thrombocytopaenia have been identified as putative indicators for severe disease [1, 24, 25]. Cerebral malaria and coma have not been associated with *P. knowlesi*, although severe malaria including hyperparasitaemia can result in cerebral pathology as shown in one reported autopsy case of *P. knowlesi* [23].

Daneshvar and colleagues reported close to 10% of hospital cases of *P. knowlesi* malaria with severe complications and a case fatality rate in their study of 1.8% [24] while others have reported higher numbers for severe cases (29% and 39% of hospital cases studied) [25, 30] and deaths (six deaths out of 22 patients with a severe case of *P. knowlesi* malaria) [30]. As of 2017, 41 deaths were reported in Malaysia [31]. Notably, deaths are being averted in recent years with improved monitoring and diagnosis and the timely use of intravenous artesunate [32] or artemisinin combination therapies (ACTs) [25, 28, 33, 34].

As this zoonotic disease remains prevalent, gaining an improved understanding of the pathogenesis of *P. knowlesi* is important. Such an understanding can be gained with the use of nonhuman primate (NHP) models [reviewed in 35, 36, 37]. Experimental infections can be designed that consider the infecting parasite species and genotype(s), the parasite stage and size of the inoculum, the duration of infections, experimental interventions, treatments, and the immune status of the host. Furthermore, tissue samples can be collected for pathology studies.

Multiple NHP species can be experimentally infected with *P. knowlesi* [reviewed in 35, 36, 38]. The most explored have been rhesus and kra monkeys. Malaria-naïve rhesus macaques will invariably succumb to *P. knowlesi* infection, unless treated with anti-malarial drugs, because of the unceasing, overwhelming rise in parasitaemia and the continued cyclical destruction of erythrocytes that occurs unabated in this species with the multiplication of the parasites every 24 hours [38–43]. Thus, rhesus macaques are viable animal models for severe disease due to hyperparasitaemia. In the course of ∼50 years, rhesus monkey infections became the main experimental host to study *P. knowlesi* biology, virulence, immune responses, antigenic variation and pathology, with much knowledge yet to be uncovered in each of these areas [e.g., see 41, 42-45] and [reviewed in 36, 37]. In stark contrast, malaria-naïve kra monkeys are able to control parasitaemia without antimalarial treatment, typically with a maximum of 1-3% parasitaemia [38-40, 46, 47], making them the preferred animal model for mild, moderate or chronic *P. knowlesi* malaria.

Despite the potential of kra monkeys for revealing mechanisms of resilience to *P. knowlesi* infections, only a few small studies have so far reported features of *P. knowlesi* infections and pathology in kra monkeys [48, 49]. Anderios and colleagues reported haematological and liver and spleen histopathology data from two kra monkeys after 60 or 90 days of infection post-inoculation either with *P. knowlesi*-infected blood from a patient or cryopreserved *P. knowlesi* (Malayan strain)-infected rhesus blood [48]. Barber and colleagues studied the deformability of RBCs from both *P. knowlesi-*infected patients and three infected kra monkeys [49]. In-depth longitudinal characterizations of clinical and other parameters during the course of *P. knowlesi* infection in this resilient host species and tissue pathology analysis will complement clinical investigations and therapeutic considerations.

By combining telemetric, clinical, parasitological and pathological data, collected longitudinally for up to 50 days, the present investigation demonstrates that kra monkeys develop and recover without treatment from clinical manifestations of malaria, including fever, thrombocytopaenia and anaemia. Low parasitaemias, low parasite tissue burdens, and overall mild tissue damage and organ dysfunction define this resilient phenotype. By highlighting the clinical and pathological consequences of *P. knowlesi*, this study provides a framework for basic and systems biological studies utilizing the *P. knowlesi* – kra monkey model for studying malaria resilience and pathogenesis (Gupta *et al*., submitted). Possible systemic mechanisms and potential research directions are discussed to better understand these phenomena and consider future novel therapies.

## MATERIALS AND METHODS

### Animal use

Experiments involving NHPs were performed at the Yerkes National Primate Research Center (YNPRC), an AAALAC International-accredited facility, following ARRIVE Guidelines and Recommendations [50]. All experimental, surgical, and necropsy procedures were approved by Emory’s IACUC and the Animal Care and Use Review Office (ACURO) of the US Department of Defense and followed accordingly. The Emory’s IACUC approval number was PROTO201700484 - YER-2003344-ENTRPR-A. 15 *M. fascicularis* (Mauritius origin, born and raised at the Mannheimer Foundation, Inc., Homestead, FL, USA), all healthy and malaria-naïve young adults (3-5 years old) were assigned within three cohorts for sequential longitudinal systems biology infection experiments, primarily aiming to understand the resilience of *M. fascicularis* to *P. knowlesi* blood-stage infections (Supplemental Table 1 and Supplemental Figs. 1-3). Male monkeys were selected to eliminate confounding anaemia measurements due to menstruation. All animals were housed socially, with 12-h light-dark cycles, in housing compliant with the Animal Welfare Act and the Guide for the Care and Use of Laboratory Animals. Environmental enrichment consisting of food and physical manipulanda was provided daily. The animals received positive reinforcement training to habituate them to ear-stick blood collections for blood smear preparations and clinical analyses requiring blood collections of less than 150 µl. The weight of the animals was determined to be between 5.6 kg and 11.7 kg at the endpoint of each experiment, when necropsies and tissue analyses were performed. The animals were anaesthetized using ketamine and euthanized via intravenous administrations of barbiturates. This euthanasia method is an acceptable method of euthanasia for NHPs per the recommendations of the “AVMA Guidelines for the Euthanasia of Animals”.

**Fig. 1.**
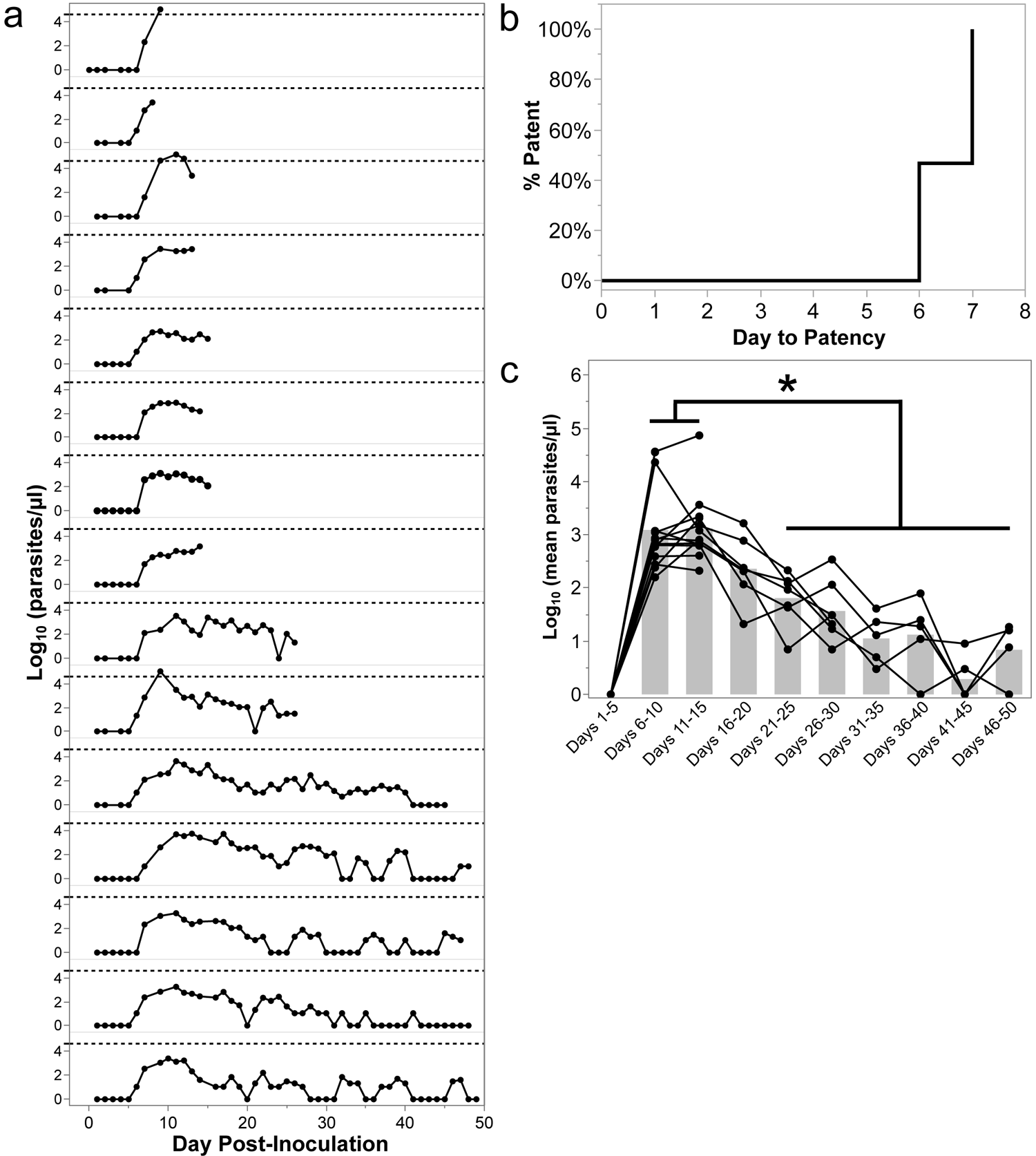
*Plasmodium knowlesi*-infected kra monkeys control parasitaemia without antimalarial treatment and develop persisting parasitaemias. a. Afternoon peripheral parasitaemias determined by thick and/or thin blood films for 15 kra monkeys infected with about 100,000 cryopreserved *P. knowlesi* H strain sporozoites. Dashed line indicates approximately 40,000 parasites/µl or about 1% parasitaemia. b. Time to patency analysis for monkeys shown in Panel a. c. Average parasitaemia for each animal during the indicated time period after infection. Light gray bars indicate mean of the shown datapoints. Statistical significance was assessed using a linear mixed-effect model followed by a Tukey-Kramer HSD post-hoc analysis. *p*-values of <0.05 were considered statistically significant. Asterisks indicate statistical significance.

### Parasite isolates and inoculations

The monkeys from experimental cohorts E07, E33 and E35 (Supplemental Table 1) were infected intravenously with ∼100,000 cryopreserved *P. knowlesi* clone Pk1(A+) sporozoites derived from the Malayan strain of *P. knowlesi* [51] (kindly provided by John W. Barnwell, Centers for Disease Control and Prevention, Atlanta, GA), which had been suspended in RPMI 1640 with 50% fetal calf serum and quick frozen, and when thawed estimated to be 0.1-1% viable; John W. Barnwell, personal communication). The Experiment 07 cohort was initially inoculated with sporozoites freshly dissected from mosquitoes at the Centers for Disease Control and Prevention, but for unexplained reasons parasitaemia did not develop in the blood. The E07 cohort was subsequently inoculated (about 80 days later) with the same batch of cryopreserved sporozoites used to infect the two other cohorts. There was no evidence that the animals had developed immunity based on similar infection kinetics across the cohorts, and significant differences were not detected in the repeated baseline transcriptomes (Gupta *et al*. submitted). Tissues from normal uninfected animals from E34, as well as archived tissue blocks from the YNPRC tissue repository, were used as negative controls for histopathological comparisons.

### Sample collections

BM aspirates from the iliac crest and venous blood collections were collected into EDTA. Capillary blood samples were collected into EDTA using standardized ear-stick procedures for CBCs and monitoring of parasitaemias.

### Telemetry

The monkeys in the E07 cohort (*n* = 7, Supplementary Table 1) had telemetry devices surgically implanted prior to infection, for real-time monitoring of temperature and other vital signs (Brady *et al*., manuscript in preparation). These devices were not included in the experimental design of the E33 and E35 protocols, which comprised two iterative studies without these measurements among the goals. In brief, the PhysioTelTM L11 telemetry implant device was secured between the external and internal abdominal oblique muscles. Raw temperature data, used in the current study, was obtained with a 1 hertz sampling frequency after which missing value indicators were removed. Hourly temperature averages were obtained using a subset of the data, minute averages obtained from the first 15 seconds of each minute if available, and then smoothed using the Hodrick-Prescott filter to reduce noise. The normal temperature range for each monkey was defined as the range of temperatures measured prior to inoculation of *P. knowlesi* sporozoites. The febrile threshold was determined by considering the parasitaemia prior to the time when the temperature rose above the individual’s threshold. The time-to-temperature response was determined as the day post-inoculation when the temperature rose above the individual’s threshold.

### Quantification of reticulocyte production index

Reticulocyte production index was calculated to assess bone marrow responsiveness to anaemia, and is given by the following equation, as described previously [52]:

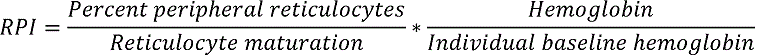

### Parasite enumeration

Parasitaemia was enumerated daily between 1 PM and 3 PM when synchronous ring-stage parasites predominated until the infections became patent. After patency, twice-daily capillary samples (8 AM, schizonts; and 1-3 PM ring stages) were acquired for parasitaemia monitoring. Thick and thin blood smears were prepared and stained with Wright’s Giemsa. Thick blood smears were used to enumerate parasitaemias below 1% by the Earle Perez method [53]. Once parasitaemia was greater than 1%, thin blood smears were used for parasitaemia measurements by determining the number of iRBCs out of 1,000 total RBCs. The number of parasites per microliter of blood was calculated from thin blood smears by multiplying the percentage of infected RBCs by the number of RBCs per microliter of blood obtained from the CBCs. Cumulative parasitaemia was calculated for each animal by adding together the daily parasitaemia (parasites/µl) from the day of inoculation to the day of necropsy. Mean parasitaemia across five-day increments was calculated for each individual by defining 5-day windows starting from the day of inoculation and averaging the parasitaemias within each window. Subsequent calculations with haemoglobin levels and platelet counts were likewise performed in this manner.

### Quantification of parasite replication rate

The replication rate of parasites was calculated as described previously [54]. Only the first peak was considered because it had the most robust sample size.

### Tissue collection, preservation, and pathology analysis

Tissues were collected at necropsy from 22 organs from all longitudinal infection experiments with the exception of E33, which had an abbreviated tissue collection, restricted to liver, lung, kidney, spleen, adrenal gland, BM, stomach, duodenum, jejunum, and colon where previous pathology had been observed. Gross lesions were photographed, and tissue sections were collected for histopathology. Each was preserved in 10% neutral buffered formalin, embedded in paraffin, sectioned at 4 μm, and stained with H&E. Diagnostic characterization and scoring of histopathology were obtained from examination of randomized, blinded H&E-stained tissues. Tissues were scored from 1 (low) to 4 (high) based on: inflammation, oedema, necrosis, haemorrhage, hyperplasia, fibrosis, and vasculitis. The scores were summed and whole organ scores obtained for comparison, as described previously with *P. vivax* infected *S. boliviensis* tissues [55].

### iRBC quantification within tissues

iRBC densities in the tissues were determined by counting the number of iRBCs in ten HPFs (1000x) under oil immersion on a standard light microscope, as described previously with *P. vivax* infected *S. boliviensis* tissues [55]. All sections were randomized and blinded.

### Erythroid progenitor measurements by flow cytometry

Five microliters of BM aspirate were placed into a 5 ml FACS tube. An antibody cocktail consisting of the antibodies outlined in Supplemental Table 9 was then added to each tube. The samples were vortexed and incubated at RT for 15 mins in the dark. The samples were washed with 500 µl of PBS followed by centrifugation at 800 × g for 7 minutes at 4°C. The supernatant was aspirated and discarded. Hoechst 33342 dye (10 ug/ml) was then added to each sample and incubated for 30 mins at 37°C in the dark. Five hundred microliters of PBS were then added to each sample followed by centrifugation at 800 × g for 7 mins at 4°C. The supernatant was then aspirated and discarded. Samples were resuspended in 100 µl of 1X Annexin V binding buffer followed by addition of the Annexin V probe. After the samples were incubated for 15 mins, an additional 400 µl of binding buffer was added, and the samples were immediately acquired on an LSR-II flow cytometer using standardized acquisition template. Voltage was controlled throughout each longitudinal experiment by calibrating the instrument using rainbow calibration particles (Biolegend). After acquisition, data were compensated in FlowJo (Treestar, Inc), and then, uploaded to Cytobank for analysis. Frequencies of erythroid cells out of the erythroid population were exported and used in analyses. Other markers in the panel were either not significantly different or are beyond the scope of this manuscript.

### Erythropoietin quantification

EPO was measured as previously described [56] with minor modifications. Frozen plasma isolated from whole blood collected in EDTA was thawed on ice. Each aliquot was then centrifuged at 750 ×*g* for 15 mins at 4°C. The supernatant was collected and used in the Quantikine IVD ELISA kit from R&D systems following the manufacturer’s suggested protocol except that the amount of suggested sample for the assays was reduced by 50% compared to what is suggested. EPO concentrations were determined using a 5-point standard curve that best fit the dynamic range of where the samples fell on the curve.

### Computation of the removal of uninfected RBCs

The degree of removal of uninfected RBCs was computed with a discrete recursive mathematical model of the RBC dynamics, as previously performed for data generated from macaques infected with *P. cynomolgi or P. coatneyi* [57, 58]. Briefly, modeling the infection trajectory of each monkey was accomplished by fitting an adaptation of the former model [58, 59] to the experimentally determined data. The model has four variables, representing reticulocytes, RBCs, iRBCs and free merozoites. The first three have a lifetime age-structure stratified into one-hour intervals, while the iRBCs are modeled with only 24 one-hour age-classes, thus enforcing the approximate 24-hour cycle of *P. knowlesi*. The model was parameterized with an RBC hazard function determined for rhesus macaques [58] using data from [60]. This hazard function allows the estimation of the level of RBC loss due to senescence. All other model estimates, reticulocyte maturation time, erythropoietic output, RBC loss due to the bystander effect, RBC loss due to parasitization and immune responses against iRBCs, are determined in a “personalized” manner, namely based on experimental data generated from each animal. The full model description can be found elsewhere [58].

### Statistical analyses

Data were generally divided into two categories for these analyses. Samples that were collected daily, namely parasitaemias, haemoglobin, reticulocyte counts, and platelet counts were divided into 5-day intervals and compared across the course of infection using a linear mixed effects model that accounted for date and monkey drop out due to sacrifice (Supplemental Figs 1-3). Direct comparisons of tissue scores were performed using a Tukey HSD test. Statistics and figures were produced in R Studio version 1.1.383, under R version 3.4.3 GUI version 1.70, or JMP Pro version 13.0.0. Associations were tested using the Spearman’s correlation test, and hierarchical multiple linear regression analyses. Multicollinearity was assessed using the olsrr package in R. Graphs were prepared using JMP Pro version 15. Comparisons were considered significant when FDR-adjusted, where appropriate, p-values below 0.05.

## RESULTS

Fifteen malaria-naïve *M. fascicularis* comprising three experimental cohorts, in addition to three control monkeys, were assigned to this study and used sequentially in a set of iterative longitudinal *P. knowlesi* infection experiments. Details about the animal cohorts (E07, E33, E34 and E35) and brief descriptions of the experimental plans are summarized in Supplemental Table 1. Schematics with parasitaemias, as well as sampling and necropsy time points for each animal by cohort are displayed in Supplemental Figures 1-3.

### Kra monkeys controlled acute *P. knowlesi* parasitaemia and developed chronic infections without anti-malarial intervention

All kra monkeys infected in this study (*n* = 15) developed patent blood-stage infections 6-7 days post-inoculation (dpi) with cryopreserved *P. knowlesi* sporozoites, and they controlled the acute phase parasitaemia without the need for anti-malarial intervention (Figs. 1a, 1b). The average *P. knowlesi* parasite replication rate between patency and peak parasitaemia was 8.59-fold ± 1.719 (mean ± SEM) per cycle, which is comparable to the value of 10-fold, which is accepted in the literature and was determined for rhesus monkey infections with *P. knowlesi* and *in vitro* culture using rhesus erythrocytes [39, 40, 62]. Most primary parasitaemic peaks averaged about 1,000 parasites/µl; however, three of the 15 monkeys developed parasitaemias that exceeded 40,000 parasites/µl, which is equivalent to about 1% iRBCs out of the total RBCs enumerated (Fig. 1a). To identify the time where the kra monkeys began controlling the parasitaemia and when chronic infections began to develop, the mean parasitaemia across five-day increments was calculated for each individual and compared. Parasitaemias peaked 6-15 dpi, declined significantly by 21-25 dpi, and began to stabilize by 31-35 dpi where they persisted at or below 200 parasites/µl (Fig. 1c). Overall, the parasitaemia kinetics in the kra monkeys was similar to those reported in previous studies irrespective of the *P. knowlesi* infections being initiated with sporozoites or blood-stages [35, 39, 40, 47]

### Kra monkeys infected with *P. knowlesi* developed mild to moderate thrombocytopaenia

Platelet levels were stable in the *P. knowlesi*-infected kra monkeys through 7 dpi, and then they began to rapidly decrease through 10 dpi while parasitaemia was rising, even though this decrease was not significantly different (Fig. 2a). After parasitaemias peaked, platelet counts began increasing and returned to baseline levels by 21-25 dpi (Figs. 2a, 2b). The monkeys experienced mild to moderate thrombocytopaenia, as reported previously for *P. cynomolgi* and *P. coatneyi* infection of rhesus monkeys [52, 60, 63], with platelet nadirs as low as 129,000 platelets/μl (Fig. 2a). Importantly, none of the animals developed severe thrombocytopaenia, defined as platelet nadirs below 50,000 platelets/µl [64]. As platelet levels decreased, the mean platelet volume (MPV) remained high but then significantly declined after about 13 dpi and remained lower than baseline throughout the rest of the infection (Fig. 2c).

**Fig. 2.**
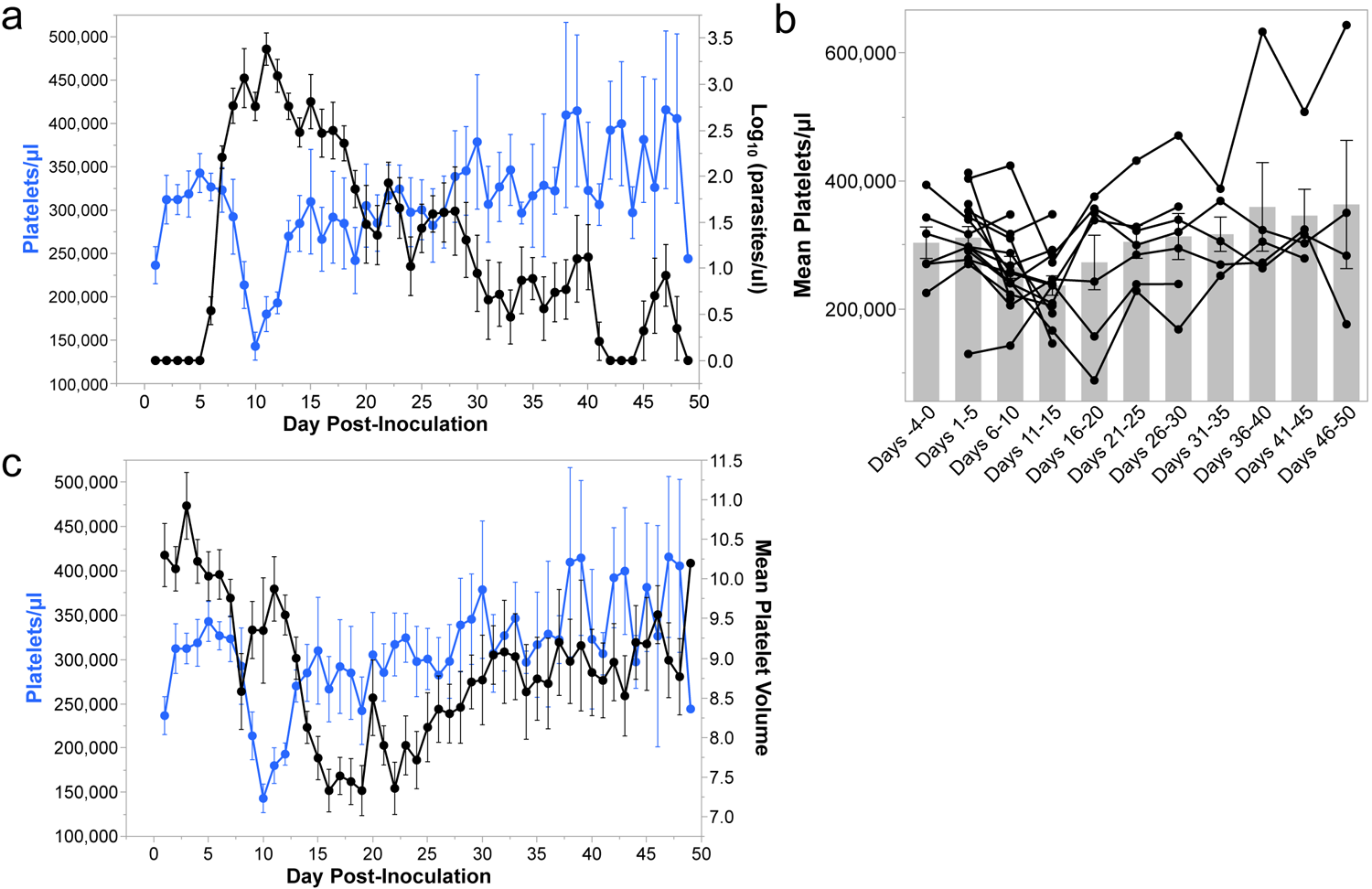
*P. knowlesi* infections in kra monkeys cause mild to moderate thrombocytopaenia. a. Platelet concentration kinetics in relation to parasitaemia before and after infection. Each dot represents the mean for all animals present during infection at that time. b. Average platelet concentration for each animal during the indicated time period after infection. Light gray bars indicate mean of the shown datapoints. c. Kinetics of platelet concentration in relation to platelet size (MPV) after infection. Statistical significance was assessed using a linear mixed-effect model followed by a Tukey-Kramer HSD post-hoc analysis. *p*-values of <0.05 were considered statistically significant. Error bars = SEM. Asterisks indicate statistical significance.

### Moderate to severe anaemia developed during *P. knowlesi* infections in kra monkeys due to removal of uninfected RBCs and not inefficient erythropoiesis

Haemoglobin levels in the infected kra monkeys were determined daily by complete blood count (CBC) analysis to evaluate the development of anaemia. Haemoglobin levels were stable through 8-10 dpi, but then sharply decreased after the peak of parasitaemia through 20 dpi (Figs. 3a, 3b). Two monkeys developed haemoglobin nadirs as low as 4.8 g/dL, corresponding to severe anaemia (< 7.0 g/dL) with about a 55% loss in haemoglobin relative to baseline (Figs. 3a, 3b). Haemoglobin levels began to increase about 21 dpi and continued to increase throughout the remainder of the infection period (Figs. 3a, 3b). Despite the dramatic loss in haemoglobin, the haemoglobin levels rebounded without anti-malarial or clinical intervention and were on average greater than 11 g/dl by 36 dpi until the end of the study (Figs. 3a, 3b). Mild anaemia persisted until the end of the experiment, and the animals never fully recovered to their pre-infection haemoglobin levels (Figs. 3a, 3b), akin to previous reports with patients [24].

**Fig. 3.**
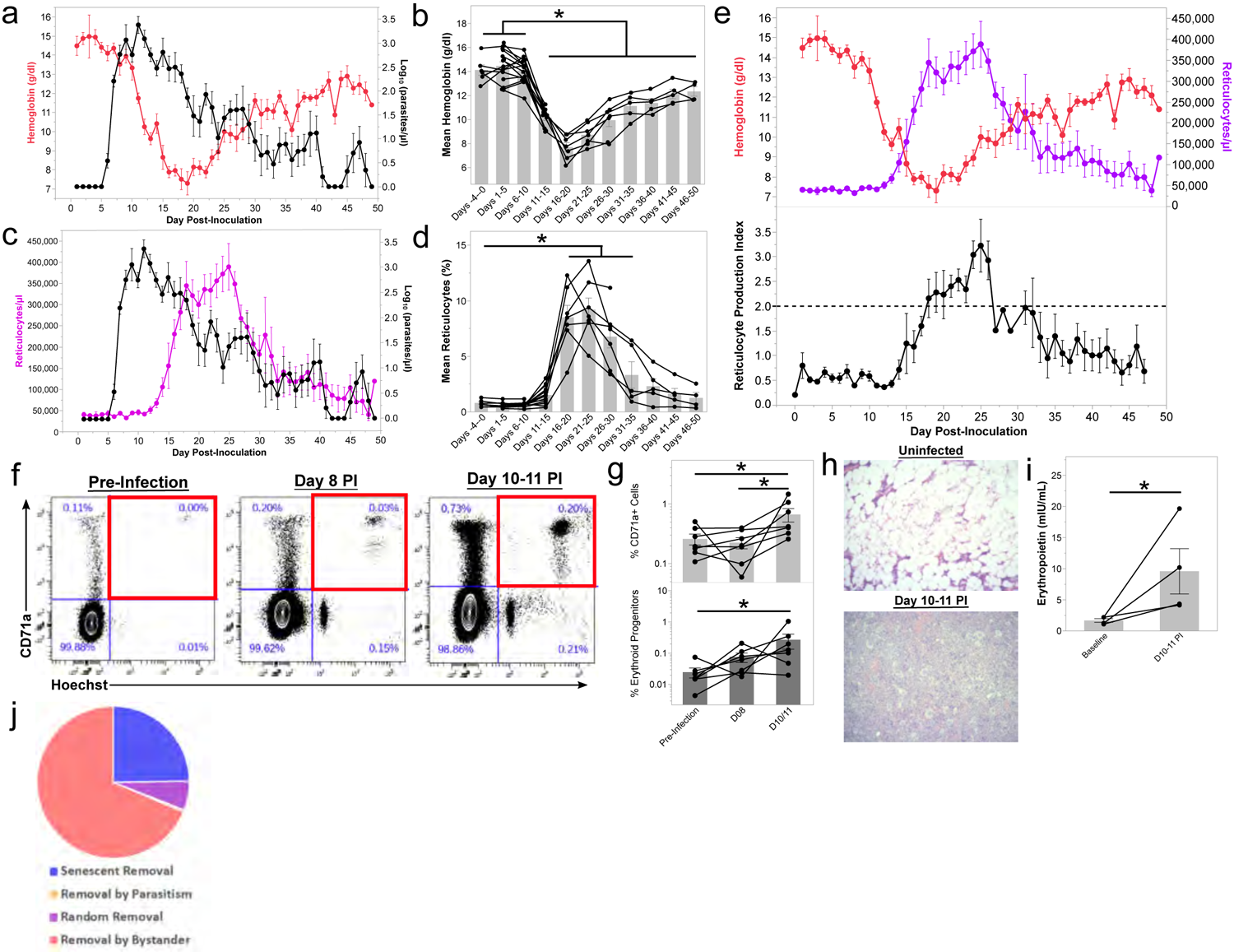
Kra monkeys experience moderate to severe anaemia after infection with *P. knowlesi* that is attributed to RBC removal rather than inefficient erythropoiesis. a. Haemoglobin concentration kinetics in relation to parasitaemia during infection. Each dot represents the mean for all animals where data were available at that time during infection. b. Average haemoglobin concentration for each animal during the indicated time period after infection. Light gray bars indicate mean of the shown datapoints. c. Platelet concentration kinetics in relation to parasitaemia before and after infection. Each dot represents the mean for all animals where data was available at that time during infection. d. Average reticulocyte concentration in peripheral blood for each animal during the indicated time period after infection. Light gray bars indicate mean of the shown datapoints. e. Relationship of haemoglobin and reticulocyte concentration kinetics with reticulocyte production index during infection. Each dot represents the mean for all animals where data were available at that time during infection. f. Representative flow cytometry plots showing the relative quantification of erythroid progenitors present in bone marrow aspirates collected before and during infection. Percentages in blue indicate the percentage of each population in the gate out of the erythroid compartment defined as SscloCD41a-CD45-LiveDead-. Gating strategy can be found in Supplementary Fig. 4. g. Relative quantification of erythroid progenitors as in Panel f. h. Representative H&E-stained bone marrow samples from uninfected, malaria and *P. knowlesi* infected kra monkeys at the indicated time points. i. Quantification of erythropoietin by ELISA in the plasma of a subset of kra monkeys after infections with *P. knowlesi*. j. Quantification of processes resulting in the elimination of RBCs using a recursive mathematical model. Statistical significance was assessed using a linear mixed-effect model followed by a Tukey-Kramer HSD post-hoc analysis. *p*-values of <0.05 were considered statistically significant. Error bars = SEM. Asterisks indicate statistical significance.

Inefficient erythropoiesis was shown to be a contributor in the development of malarial anaemia in rhesus monkeys with *P. coatneyi* or *P. cynomolgi* infections [56, 60], as well as in humans and rodents infected with malaria parasites [65–68]. Thus, it was hypothesized that inefficient erythropoiesis may contribute, at least in part, to the development of anaemia in *P. knowlesi* infected kra monkeys. Contrary to this hypothesis, however, peripheral reticulocyte counts steadily increased as haemoglobin levels decreased, and they peaked a couple of days after the haemoglobin levels reached their nadirs (Figs. 3c-e). To evaluate if the increase in reticulocyte levels was appropriate to compensate for the decrease in haemoglobin levels, the reticulocyte production index (RPI) was calculated. Indeed, the RPI was sustained over 2 from about 17-26 dpi, indicating that there was adequate compensation by the bone marrow (BM), following the dropping haemoglobin levels, and no evidence of inefficient erythropoiesis was found (Fig. 3e).

To confirm if the increase in peripheral reticulocyte levels was due to upregulation of erythropoiesis in the BM, the frequency of erythroid lineage cells in the whole BM aspirates collected prior to infection and at 8-11 dpi was evaluated by flow cytometry. The gating strategy used to quantify the frequency of erythroid progenitor cells in the BM aspirates is described in Supplemental Fig. 4. The erythroid lineage cells were defined as CD41a-CD45-LiveDead-CD71+Hoechst+ cells. The frequency of erythroid lineage cells was increased in the BM aspirates, acquired by 10 or 11 dpi, compared to baseline (Figs. 3f-g). Consistent with the flow cytometry data, haematoxylin and eosin (H&E)-stained BM specimens acquired at necropsies as early as 10 dpi revealed substantial expansion of erythroid progenitors relative to uninfected controls (Fig. 3h). Correspondingly, plasma erythropoietin (EPO) levels were significantly elevated above baseline levels at these times for a subset of animals where plasma samples were available (Fig. 3i).

**Fig. 4.**
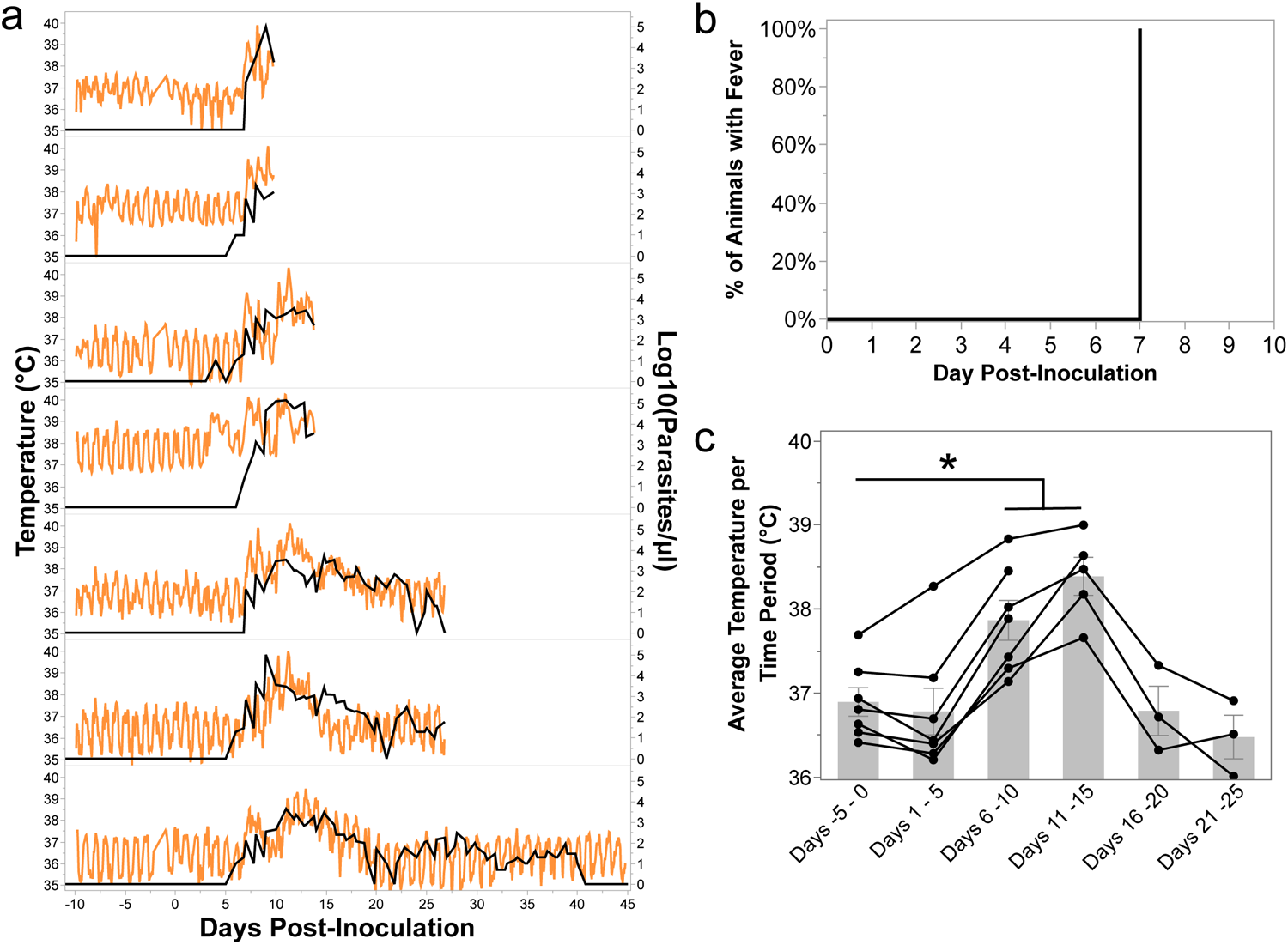
Kra monkeys develop fevers shortly after patency, which resolve after the control of parasitaemia. a. Temperature kinetics in relation to parasitaemia in animals that had telemetry devices surgically implanted. b. Time to fever analysis using the telemetry data in panel a. c. Average temperature for each animal during the indicated time period after infection. Light gray bars indicate mean of the shown datapoints. Error bars = SEM. Bar graphs and error bars represent mean ± SEM. Statistical significance was assessed using a linear mixed-effect model followed by a Tukey-Kramer HSD post-hoc analysis. *p*-values of <0.05 were considered statistically significant. Asterisks indicate statistical significance.

Since there was no evidence of inefficient erythropoiesis in the *P. knowlesi*-infected kra monkeys and the relatively low parasitaemia could not account for the observed decrease in haemoglobin levels, the degree of removal of uninfected RBCs was computed using a mathematical model devised for this purpose [58]. Modeling of the kra monkey data showed an appropriate timely compensatory erythropoietic response, with 68.9% ± 4.9% (mean ± SEM) attributable to the removal of uninfected bystander RBCs (Fig. 3j). The model estimated that the parasite was responsible for less than 0.5% of the total RBC losses (Supplemental Fig. 5).

**Fig. 5.**
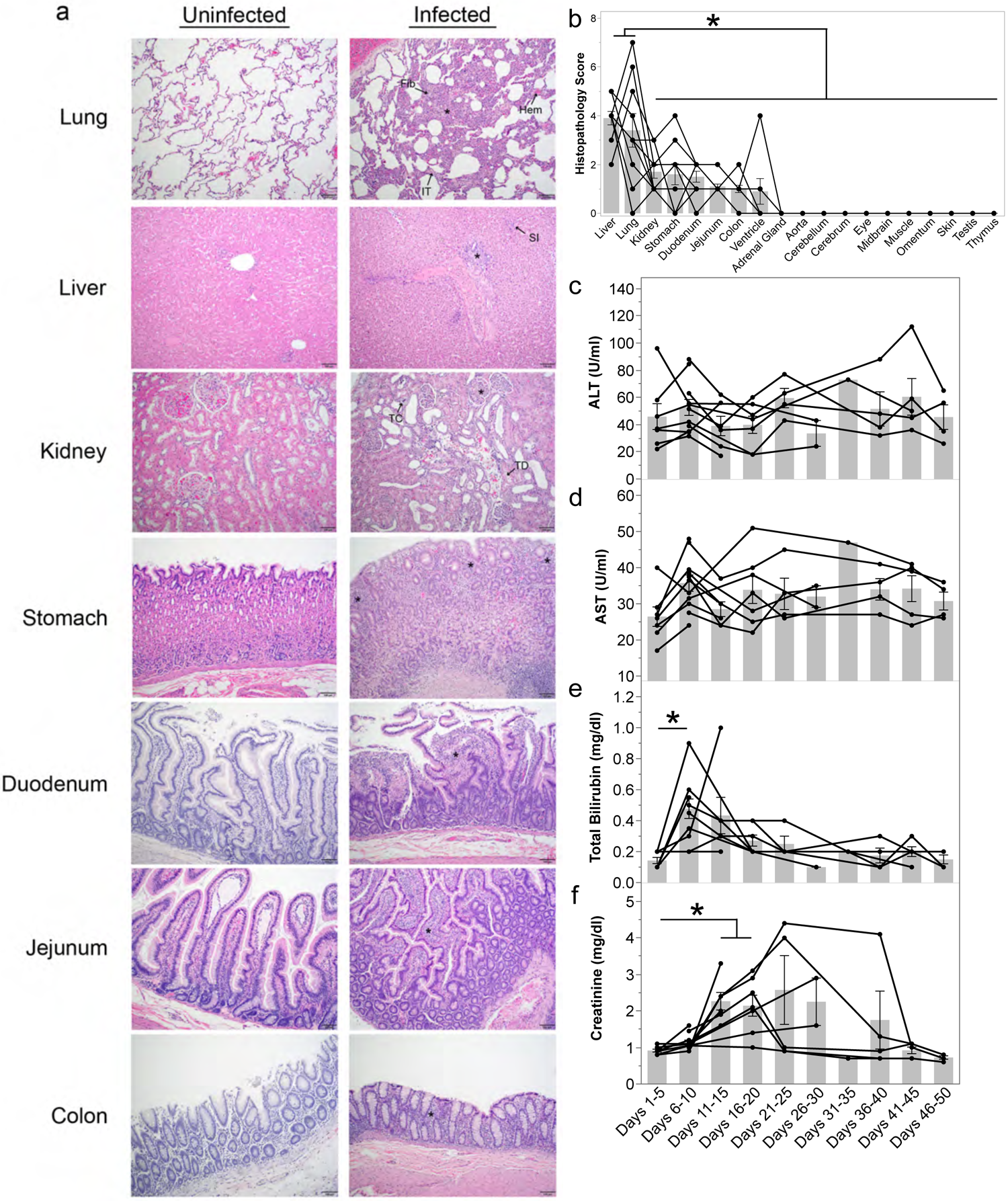
*Plasmodium knowlesi*-infected kra monkeys exhibit histopathology consistent with systemic measures of disease. a. Representative light micrographs of H&E-stained tissue sections from control and infected kra monkeys. Infected lungs showed areas of fibrosis (Fib), hyperplasia (*), haemolysis (Hem), and interstitial thickening (IT). All animals had periportal (*) and sinusoidal (SI) infiltration in the liver. Kidney histopathology included tubular crystal formation (TC), hypercellular glomeruli (*), and tubular degeneration (TD). The stomach, duodenum, jejunum, and colon all exhibited mucosal immune infiltration (*). b. Semi-quantitative histopathology scores of tissues collected at necropsy. The average alanine transaminase (c), aspartate aminotransferase (d), total bilirubin (e), and creatinine (f) for each animal where data were available during the indicated time period after infection. Light gray bars indicate mean of the shown datapoints. Error bars = SEM. Statistical significance was assessed using a linear mixed-effect model followed by a Tukey-Kramer HSD post-hoc analysis. *p*-values of <0.05 were considered statistically significant. Asterisks indicate statistical significance.

### Kra monkeys infected with *P. knowlesi* developed fevers that self-resolved

Densely sampled temperature data were analyzed from the kra monkeys that had surgically implanted telemetry devices (*n* = 7, Supplemental Table 1) to study their temperature patterns including possible fever after infection (Fig. 4a). Fever was defined as any increase above an individual’s baseline range of temperatures, which represent each monkey’s daily circadian rhythm. All kra monkeys exhibited a fever by 7 dpi corresponding roughly to the period in which the blood-stage infection reached patency, between 6-8 dpi (Figs. 4a, 4b). The average temperature of the animals significantly increased from pre-infection values of 36.89 ± 0.17 °C (mean ± SEM) to 37.87 ± 0.24°C and 38.39 ± 0.23°C by 6-10 and 11-15 dpi, respectively (Fig. 4c). The febrile threshold [62] was defined as the parasitaemia at which the kra monkeys developed temperatures and determined to be about 160 ± 71.58 parasites/μl (mean ± SEM). After parasitaemia was controlled, the average temperatures decreased to pre-infection levels of 36.79 ± 0.29 °C and 36.48 ± 0.26 °C for the day 16-20 and 21-25 time periods, respectively (Figs. 4a and 4c). Monkeys that proceeded to establish chronic low-level parasitaemia no longer experienced temperature spikes (Figs. 4a, 4c).

### *P. knowlesi* infection in kra monkeys caused mild to moderate tissue pathology

An extensive analysis was performed of H&E-stained tissue sections acquired from up to 22 different organs collected from the *P. knowlesi*-infected kra monkeys, sacrificed at different dpi, to determine if and the extent to which the monkeys may have sustained tissue damage. Hyperplasia, interstitial thickening, and fibrosis were observed in the lungs of nearly all monkeys (13 / 15), and mild to moderate haemorrhage occurred in four of them (Fig. 5a and Supplemental Table 2). All monkeys had Kupffer cell hyperplasia and evidence of liver inflammation based on cellular infiltrates around the periportal region and congestion in the hepatic sinuses (Fig. 5a). No hepatocellular necrosis or cholestasis was noted. The kidneys of all animals showed glomerular hypercellularity without glomerulonephritis, and three animals had tubular degeneration (Fig. 5a and Supplemental Table 2). One monkey with high parasitaemia and elevated creatinine had evidence of calcification or crystalline deposition in the kidney tubules (Fig. 5a and Supplemental Table 2). Less common kidney pathologies included interstitial inflammation and haemorrhage (Supplemental Table 2). Gastrointestinal inflammation was observed in the tissues from each monkey and included gastritis, duodenitis, jejunitis, and mild colitis without oedema. The spleens from the infected animals were grossly enlarged and weighed significantly more than the spleens from uninfected animals; the day of necropsy did not influence these results (Supplemental Table 3). The adrenal glands, aorta, cerebrum, cerebellum, midbrain, eyes, lymph nodes, omentum, skeletal muscle, testis, and thymus were unremarkable.

The severity of the observed tissue damage in each tissue except for the BM and spleen was scored for each animal to facilitate a semi-quantitative comparison of tissue damage between organs (Supplemental Table 2) as performed previously with tissues from *P. vivax* infected *Saimiri boliviensis* monkeys [55]. The severity of tissue damage in the lungs and liver was significantly higher than other tissues that sustained pathology (Fig. 5b and Supplemental Table 4). The date of necropsy – which differed for the animals – did not affect the histopathological scores, indicating that how long an animal was parasitaemic did not have a detectable impact on the severity of tissue damage (Supplemental Table 5). In line with the absence of cholestasis and hepatocellular necrosis, aspartate aminotransaminase (AST) and alanine aminotransferase (ALT) remained unchanged during infection, and while bilirubinaemia was observed, it resolved by 11-15 dpi (Fig. 5e). Consistent with kidney pathology, serum creatinine levels were significantly elevated 11-20 dpi before returning to baseline levels (Fig. 5f). Notably, creatinine levels varied significantly between individuals after 20 dpi (Fig. 5f).

### Parasite accumulation in the vasculature is associated with severity of tissue damage

To determine if the severity of damage in each tissue was associated with parasite accumulation within that tissue, the number of infected RBCs (iRBCs) in 10 high-power fields (HPFs) was enumerated (Supplemental Table 6). This analysis provided approximate iRBC densities within each tissue, for 22 tissue types, as also performed previously with *P. vivax* infected *S. boliviensis* tissues [55]. A representative image of parasites identified in tissues is shown in Fig. 6a. Multiple analyses were then used to assess if the accumulation of parasites in the tissues was associated with the tissue damage scores (Supplemental Table 2). First, a univariate analysis was performed between tissue scores and iRBC density. Parasite iRBC burden weakly positively correlated with tissue score (ρ = 0.265, p-value = 0.00022). Next, a hierarchical multiple linear regression analysis was performed to determine which factors affected tissue scores the most (Supplemental Table 7). A model considering only iRBC tissue burden did not result in a linear relationship (adjusted R^2^ = 0.0354, p-value = 0.0054). However, when combined with the tissue type, the relationship was moderately linear (adjusted R^2^ = 0.678, p-value = 2.2 x 10^-16^), indicating that the iRBC burden in a tissue may contribute to the degree of damage observed. The day of necropsy, parasitaemia at necropsy, and cumulative parasitaemia were also tested and were found not to contribute to the relationship (Supplemental Table 8). No multicollinearity was noted.

**Fig. 6.**
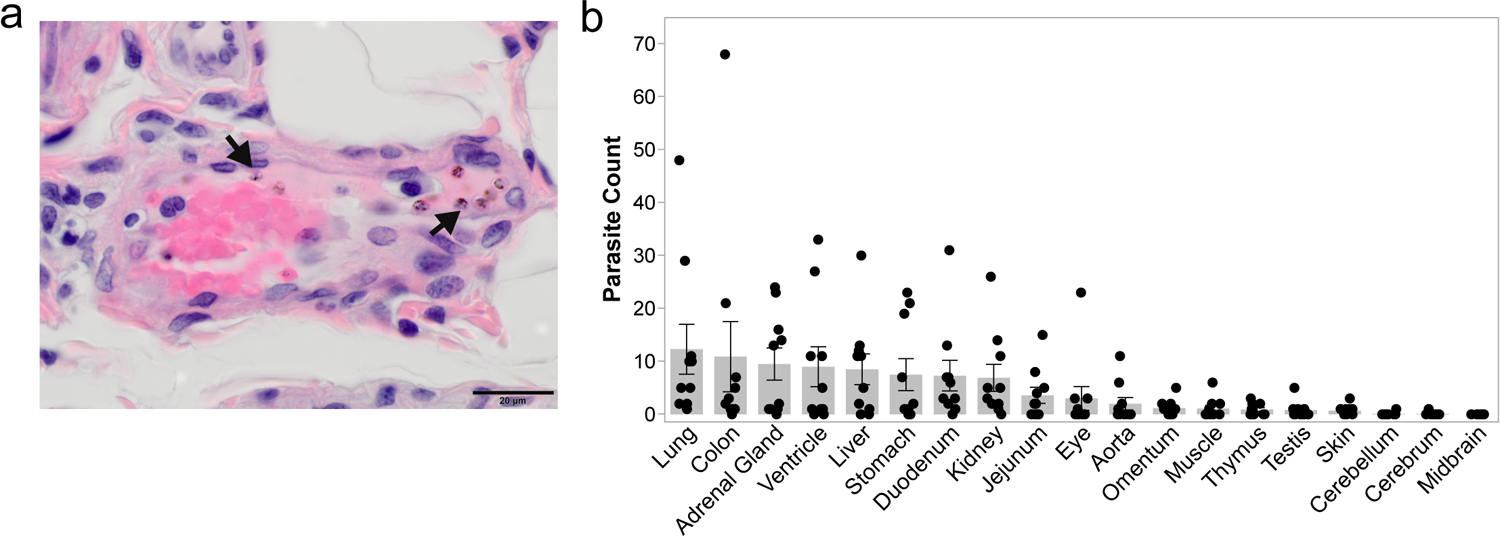
Parasites accumulate in the tissues of *P. knowlesi*-infected kra monkeys. a. Representative micrograph showing parasites (arrows) in H&E-stained gastrointestinal submucosa. b. Average parasite counts in 10 high power fields of 22 different tissues from *P. knowlesi*-infected kra monkeys.

## DISCUSSION

This longitudinal infection study defines clinical, physiological and pathological changes that occurred in kra monkeys after being experimentally infected with *P. knowlesi* sporozoites. Consistent with kra monkeys being a resilient host, they controlled their parasitaemia soon after becoming patent and did not develop severe thrombocytopaenia, a complication that has been observed with severe human *Plasmodium* infections [reviewed in 64], and most notably with *P. knowlesi* [1, 24, 25], as well as some *P. cynomolgi* or *P. coatneyi* infections in rhesus monkeys [52, 60]. Recently, thrombocytopaenia has been associated with platelet-*Plasmodium* iRBC interactions that result in the killing of those infected cells [69]. While platelet killing of *P. knowlesi*-iRBCs was not evaluated in the current study, it is certainly conceivable that platelets could have a killing function in this animal model. In fact, the mild thrombocytopaenia observed here would be fitting with platelets having a role in eliminating parasites and contributing to the lower level of iRBCs in this resilient host. It would be of extreme interest to test this possibility and establish direct comparisons of parasitaemia and thrombocytopaenia with platelet killing assays also performed during rhesus infections, where the parasitaemia escalates unabated to high levels.

A precipitous and dramatic drop of haemoglobin occurred during the peak of *P. knowlesi* parasitaemia in the kra monkeys and continued until the parasitaemia was controlled. This resulted in haemoglobin nadirs within the documented range of moderate to severe malarial anaemia experienced by *P. cynomolgi* or *P. coatneyi*-infected rhesus macaques [52, 60, 63, 70], humans with *Plasmodium* parasites [reviewed in 71], and notably children with *P. knowlesi* [26]. These haemoglobin kinetics are reproducible in NHPs infected with *Plasmodium* spp. and virtually indistinguishable from the haemoglobin kinetic data generated retrospectively from neurosyphilitic patients infected with *Plasmodium vivax* [72]. The development of severe anaemia in kra monkeys infected with *P. knowlesi* has not been previously reported and these data raise the intriguing point that these animals demonstrated natural resilience to their infection despite demonstrating this disease manifestation. This raises questions regarding what mechanisms are used by these animals to overcome the anaemia.

Interestingly, in this regard, in contrast to inefficient erythropoiesis reported for humans, rhesus macaques, and rodents infected with malaria parasites [56, 60, 63, 65–68, 70], there was no evidence in the *P. knowlesi*-infected kra monkeys that inefficient erythropoiesis contributed to the development of their anaemia. Instead, the kra monkey BM responded appropriately when EPO was elevated in the plasma, comparable to data shown recently for four kra monkeys infected with *P. coatneyi*-infected RBCs and monitored for anaemia [70]. Thus, the mechanism(s) that apparently limits an EPO-dependent response to malarial anaemia in humans, rhesus macaques, and rodent malaria parasite model systems, appears to be absent in kra monkeys, thereby, enabling kra monkeys to sufficiently compensate for the loss of RBCs during *P. knowlesi* malaria. The data presented here suggest that preserving the BM’s ability to compensate for the loss of RBCs is a noteworthy characteristic of resilience displayed by the kra monkeys to *P. knowlesi* infections. Future comparative studies should explore soluble factors such as inflammatory molecules, metabolites, growth factors, etc., that are different between kra and rhesus monkeys infected with *P. knowlesi* to identify factors that may contribute to inefficient erythropoiesis in the rhesus but not the kra monkey.

Previous malarial anaemia studies based upon rhesus macaques infected with *P. cynomolgi* or *P. coatneyi* could not readily distinguish the contribution of inefficient erythropoiesis versus removal of uninfected erythrocytes because these processes occurred sequentially [52, 60, 63, 70]. In rhesus macaques infected with *P. cynomolgi* or *P. coatneyi*, there is apparent inefficient erythropoiesis that occurs while haemoglobin levels begin to drop [52, 60, 63, 70]. This is followed by a rapid decline in haemoglobin levels despite a compensatory response by the BM [52, 56, 60, 63, 70]. Mathematical modelling of RBC haemodynamics in rhesus monkeys infected with *P. coatneyi* suggested that removal of uninfected erythrocytes contributed more to anaemia in the *P. coatneyi* infected rhesus monkeys than inefficient erythropoiesis [57]. Significant loss of uninfected RBCs was also predicted based on the retrospective modeling of *P. falciparum* parasitaemia and anaemia data from neurosyphilis patients undergoing malariotherapy [73] and demonstrated in clinical studies involving *P. falciparum* and *P. vivax* [74]. Still, the extent that inefficient erythropoiesis contributed to the exacerbation of anaemia in the animal models, dominated by the removal of uninfected RBCs, remained unclear. Curiously, the kra monkeys in this study displayed no evidence of inefficient erythropoiesis yet maintained similar adverse haemoglobin kinetics as determined from rhesus monkey models [52, 56, 60, 63, 70] and human data [reviewed in 71]. In kra monkeys infected with *P. coatneyi,* the observed erythropoietic response was suboptimal, and alone did not explain the anaemia observed in this species [70]. Overall, with malaria caused by various *Plasmodium* species, it appears that the removal of uninfected RBCs may contribute more than inefficient erythropoiesis to the development of malarial anaemia than previously appreciated, as also generally concluded by Jakeman and colleagues [73].

Kra monkeys infected with *P. knowlesi* are well-suited to study the possible physiological and immunobiological mechanisms that lead to the removal of uninfected RBCs, whether in malaria-naïve or semi-immune animals [60]. Interestingly, recent RBC deformability studies showed that the ability of uninfected RBCs to become deformable was not affected in *M. fascicularis* (*n* = 3, bred and grown in animal facilities at Nafovanny in a malaria-free environment in Vietnam) after being infected for 8 days with *P. knowlesi* (UM01 strain) blood-stage parasites, and echinocyte formation was not observed [49]. These findings contrast with that group’s complementary results from human infections [49], and published findings from infected rhesus monkeys [75, 76], in support of the idea that the absence of these adverse rheologic features and the maintained deformability of RBCs in the kra monkeys supports their resilience. Future studies are warranted that look more closely at all of these factors side-by-side with both monkey species, to hone in on the various mechanisms that may support the resilient phenotype.

Acute *Plasmodium* infections caused by different parasite species – whether in NHPs or humans – are known to cause damage to tissues and vital organs and contribute to clinical complications and possibly death [reviewed in 32]. Anderios and colleagues described histological changes in the spleen and liver of two kra monkeys (born and bred at the Institute for Medical research in Kuala Lumpur) after 60 or 90 days of infection with blood-stage parasites derived from the blood of a human patient or a cryopreserved isolate from an infected rhesus (obtained from the American Type Culture Collection (ATCC)) [48]. The current longitudinal study with 15 kra monkeys is much more comprehensive with the histopathological analysis of 22 tissues. Here, histological changes in the liver, lungs, kidneys, and the gastrointestinal tract were identified, but consistent with kra monkeys being a resilient host, the tissue lesions observed were relatively mild (Supplementary Table 2). These results can be contrasted with the severe tissue damage previously reported from *P. knowlesi*, *P. coatneyi* or *P. cynomolgi* infections in rhesus macaques [45, 77–80].

Consistent with the mild histological changes observed in this study of kra monkeys, there were only a few indications of organ dysfunction. Nearly all animals had some degree of tissue damage in the lungs, and the lungs had the highest iRBC count per high power field. Creatinine levels became elevated at the peak of parasitaemia and remained elevated, indicative of kidney issues. Given the minor tissue-damage observed in the kidneys, it is possible that the increased creatinine could be related to an increase in antigen-antibody immune complex deposition in the kidney tubules or a decrease in fluid intake. Regardless, it is relevant to note that increased creatinine has been a common finding in patients with severe malaria [29]. Hyperbilirubinaemia has also been associated with severe *P. knowlesi* malaria in humans [24]. Likewise, bilirubin was elevated in the kra monkeys. Although direct and indirect bilirubin levels were unavailable, the absence of significant histological evidence for hepatic injury such as cholestasis suggests that the elevated total bilirubin levels in the kra monkeys were due to hemolysis of uninfected RBCs. In agreement with this hypothesis, bilirubinaemia returned to baseline in the latter part of the infections, coincident with the recovery of haemoglobin levels. Neither ALT nor AST were elevated at any stage in the infections, which is consistent with the absence of hepatocyte necrosis, or signs of major tissue damage systemically. Together, these data suggest that kra monkey resilience to *P. knowlesi* infection, as demonstrated by their control of parasitaemia and compensation for anaemia, is reflected by only minimal or mild tissue damage and organ dysfunction.

The presence of iRBCs in various tissues have been reported from previous studies with *P. knowlesi* infected rhesus macaques [44, 45]. Likewise, *P. knowlesi* iRBCs were identified in the tissues of the kra monkeys in this study and they predominated in the spleen, adrenal glands, lungs, kidneys, and gastrointestinal tract. However, relatively few iRBCs were identified, which is consistent with the comparatively low number of circulating parasites in the resilient kra monkey species. Regardless, similar as determined in prior studies with *P. vivax* infection of *Saimiri boliviensis* monkeys [55], there was a modest relationship in the current study between histopathology score and parasite accumulation in the tissues.

Understanding the immune responses and pathophysiological processes that occur during *P. knowlesi* infections in humans has been of great interest but studying these infections longitudinally in humans is not feasible due to ethical requirements to treat individuals once they are parasitaemic. *Plasmodium knowlesi*-infected kra monkeys are a viable alternative for studying the progression of acute and chronic infections, and without drug treatment being a confounding factor. Unlike infections of rhesus monkeys with various *Plasmodium* species, the *P. knowlesi*-infected kra monkeys uniformly develop naturally occurring chronic infections [39, 40, 46, 47], and with similarly low parasitaemia levels as recorded for uncomplicated *P. knowlesi* infections in humans [24]. Importantly, while the kra monkeys demonstrated resilience from severe and deadly disease, chronicity was characterized similar to human infections by persistent and lower than normal haemoglobin levels. These results warrant further exploration in the context of this animal model to understand the loss of the uninfected RBCs and to determine what host responses may be occurring.

Interestingly, *P. knowlesi* infections in humans induce fevers at lower parasitaemias than *P. falciparum* infections [25]. Given recent findings that macaque RBC culture-adapted *P. knowlesi* parasites could be adapted to human RBCs by acquisition of sialic acid-independent invasion mechanisms, one hypothesis to explain this is that *P. knowlesi* is not as well-adapted to humans, and, thus, can trigger immune responses more quickly than *P. falciparum* [81]. Therefore, it was expected that kra monkeys, as a natural host of *P. knowlesi,* would develop fevers once high parasitaemias developed. However, the kra monkeys developed fevers shortly after their infections reached patency, and their pyrogenic thresholds were about 160 parasites/µl. These data suggest that the lower pyrogenic threshold observed for *P. knowlesi* in humans may not solely be due to poor adaptation to humans. Kra monkeys may have evolved adaptations in pattern recognition receptors that enable them to detect parasite-derived molecules at lower levels than other primates, thereby, enabling the immune system to detect and respond to *P. knowlesi* more quickly, as shown in one study with a few animals [46].

As this research field advances, delving deeper into specific host-parasite interactions and pathways, it is important to take into consideration the origin and genetic background of the host monkeys and the infecting parasites, as well as prior experimental history of the animals and whether experiments are initiated with sporozoites or blood-stage parasites. The source and validation of the monkey and parasite species are critical. Thus, we have taken care to indicate the macaque monkey and *P. knowlesi* parasite sources and isolate information in this manuscript, and to the extent available for referenced papers [48, 49]. As early as 1932, major differences and nuances of *P. knowlesi* parasite-host combinations were described by Napier and Campbell [82] and Knowles and Das Gupta [38], and such data summarized then from numerous NHP infections, and since by others [39, 40, 83, 84], including instances of ‘hybrid’ short-tailed kra/cynomolgus-rhesus monkeys [83], may bring important if not critical insights and value to current research.

## CONCLUSIONS

This study reports a systematic analysis of *P. knowlesi*-infected kra monkeys to identify basic features of resilience to malaria, which could be relevant for infections caused by multiple species of *Plasmodium*. This investigation affirms previous reports that kra monkeys are resilient to *P. knowlesi* infections and shows that their resilience is not – at least in part – because they completely subvert development of clinical signs of malaria and avoid tissue-damage. Instead, while kra monkeys prevent *P. knowlesi* parasitaemia from rising above 1-3% they mount compensatory physiological responses that may reverse disease progression and limit tissue pathology. The kra monkey-*Plasmodium* infection model system is valuable for studying mechanisms of resilience to malaria and identifying specific physiological and immunobiological responses that may function to minimize disease and death in people infected with *Plasmodium*. Of note, the kra monkey data best reflected the clinical picture of children rather than adults infected with *P. knowlesi,* as determined in a few studies [26–28]. Continued validation of this animal model is warranted as a complement to human studies aimed at understanding mechanisms of resilience to malaria, with the possible goal of new host-directed therapies to resolve acute disease states and the progression of chronic *Plasmodium* infections. Such inquiry has begun, specifically, for example, involving in-depth analysis of the peripheral blood transcriptomes of rhesus and kra monkeys in response to *P. knowlesi* (Gupta *et al*., submitted).

## SUPPLEMENTAL TABLE LEGENDS

**Supplemental Table 1.** Macaque Cohort and Experimental Summaries. Details regarding four monkey cohorts involved in the current study are summarized; three of these were experimentally infected with *P. knowlesi* sporozoites, and one served as control group. The cohorts are listed in the order in which experiments using these animals were performed. These animals and their longitudinal infection designs were part of a systems biology program, with iterative cohort experimentation designed to satisfy the goals of those research programs. The non-sequential experimental numbering (E07, E33, E34, and E35) reflects the experimental numbers assigned in the MaHPIC Laboratory Information Management System. Telemetry data were collected for temperature, blood pressure, heart rate and activity level (manuscript in preparation). Spx is an abbreviation for splenectomy.

**Supplemental Table 2.** Summary of histopathology scores for *P. knowlesi*-infected macaques. Semi-quantitative scores are presented as a heatmap to visually represent severity, with 0 being no changes or damage, and 4 being diffuse changes. Organs were scored in several categories, including inflammation, oedema, crypt inflammation (for the gastrointestinal organs), necrosis, haemorrhage, hyperplasia (including alveolar wall thickening/interstitial hyperplasia, for the lungs, Kupffer cell hyperplasia in the liver, and glomerular hyperplasia for the kidneys), fibrosis, vasculitis, and tubular degeneration (for kidneys). The histopathology categories shown reflect the range of pathologies noted, whether for one or multiple monkeys.

**Supplemental Table 3.** Linear regression: spleen weight *vs*. time of infection. The effect of time at necropsy on spleen weight was tested via a linear regression model and found not to contribute.

**Supplemental Table 4.** Tukey HSD Post-hoc pairwise comparison of tissue score. Semiquantitative tissue scores were compared pairwise using Tukey HSD post-hoc analysis. Mean difference is the difference in means between the organ pair being compared. Adjusted *p*-value, significance and upper and lower bound are included. \**p* <0.05; \*\**p* < 0.005, \*\*\**p* <0.0005, \*\*\*\**p* <0.00005; NS = not significant.

**Supplemental Table 5.** Linear regression: pathology score *vs*. necropsy day. The effect of time at necropsy on pathology score was tested via a linear regression model and found not to contribute.

**Supplemental Table 6.** Summary statistics for parasite tissue counts. Parasites in H&E-stained tissue sections for each organ were quantified by light microscopy. Mean number of parasites in 10 high power fields per H&E-stained tissue, standard deviation, standard error, and number of sections are presented.

**Supplemental Table 7.** Hierarchical Linear Regression Analysis. Multiple Linear Regression (MLR) was performed to test the relationship between score, count, and organ. For organ, adrenal gland was selected as the reference tissue. Parameter is significant at α = 0.05; \**p* < 0.05, \*\**p*<0.005, \*\*\**p*<0.0005.; NS= not significant.

**Supplemental Table 8.** Hierarchical Linear Regression Analysis of Infection Parameters. Multiple Linear Regression (MLR) was performed to test the relationship between score, count, days post-inoculation (DPI), parasitaemia, and cumulative parasitaemia. Parameter is significant at α = 0.05; \**p* < 0.05, \*\**p*<0.005, \*\*\**p*<0.0005.; NS= not significant.

**Supplemental Table 9.** Flow cytometry staining cocktail for measuring erythroid progenitors in rhesus and kra monkey bone marrow aspirates.

## SUPPLEMENTAL FIGURE LEGENDS

**Supplemental Fig. 1.** E07 Experimental Design and Parasitaemia. Pilot *P. knowlesi* infection in 7 kra monkeys (11C131, 11C166, 12C36, 12C44, 12C53, H12C59, H12C8) with staggered necropsy endpoints. Top Panel: Schematic of the planned (generalized) experimental design with pre-infection surgical implantation of a telemetry device (scissors) plus recovery time and their activation for collection of physiological data, before (grey bar) and after surgery baseline timepoint (TP) sample collections (gold bars), *P. knowlesi* cryopreserved sporozoite inoculations at day 0 (syringe), predicted parasitaemia kinetics (pink curved line) with early infection, log-phase, peaking parasitaemia, and sequential chronic phase TPs indicated for blood and bone marrow sample collections. Necropsy endpoints (*) were planned for selected animals at the acute and both early and late chronic stages of infection. Bottom Panel: Schematic showing E07 experimental data including *P. knowlesi* cryopreserved sporozoite inoculations on day 0 (syringe^†^), daily parasitaemias graphed (pink line), and defined TPs (gold bars) and the specific days of euthanasia and necropsy endpoints (*) are indicated for each of the animals. No subcurative treatments were required, as the blood-stage infections and clinical signs naturally began to resolve, as expected with kra monkeys. ^†^This cohort had previously been inoculated with sporozoites freshly isolated from mosquito salivary glands, yet for reasons unknown, blood-stage parasitaemia did not result. To what extent the immune system may have been stimulated at the time, or not, remains an open question.

This pilot experiment allowed for 1) testing of cryopreserved stocks of *P. knowlesi* sporozoites and the kinetics of the parasitaemia in kra monkeys (pink line); 2) capturing blood, bone marrow and tissue data on both the acute and chronic phases of the infection; 3) testing of experimental and analysis pipelines for the generation and analysis of biological data at specific TPs, including the secure and reliable transfer of samples or data as required across the large MaHPIC and HAMMER consortia [85]; and collection and analysis of infected tissue samples from necropsies.

**Supplemental Fig. 2.** E33 Experimental Design and Parasitaemia (Iterative *P. knowlesi* infections in a cohort of 4 kra (13C90, 14C15, 14C3, H13C110) monkeys to study acute infections and also establish and study chronic infections. Top Panel: Schematic of the planned (generalized) experimental design with timepoints (TP) of sample collection (gold bars), *P. knowlesi* cryopreserved sporozoite inoculations at day 0 (syringe), predicted parasitaemia kinetics (pink line) with early infection, log-phases, peaking parasitaemia, and later TPs indicated for blood and bone marrow sample collections. Necropsy endpoints (*) were planned for each species after day 14, prior to the natural decline in parasitaemia as observed for kra monkeys in E07 (Supplemental Fig. 1). Bottom Panel: Schematic showing E33 experimental data including *P. knowlesi* cryopreserved sporozoite inoculations on day 0 (syringe), daily parasitaemias graphed (pink lines), and defined TPs (gold bars) and the specific days of euthanasia and necropsy endpoints (*) are indicated for each of the animals. No treatment was required.

**Supplemental Fig. 3.** E35 Experimental Design and Parasitaemia. Iterative *P. knowlesi* infection in a cohort of 4 kra monkeys (13C33, 13C74, H13C101, H14C17) to study acute and chronic infections. Top Panel: Schematic of the planned (generalized) experimental design with timepoint (TP) sample collections (gold bars), *P. knowlesi* cryopreserved sporozoite inoculations at day 0 (syringe), predicted parasitaemia kinetics (pink line) with early infection, log-phases, peaking parasitaemia, and later TPs indicated for blood and bone marrow sample collections. Necropsy endpoints (*) were planned for each species after day 14, prior to the natural decline in parasitaemia as observed for kra monkeys in E07 (Supplemental Fig. 1), and when parasitaemia was still undetectable. Bottom Panel: Schematic showing E35 experimental data including *P. knowlesi* cryopreserved sporozoite inoculations on day 0 (syringe), daily parasitaemias graphed (pink lines), and defined TPs (gold bars) and the specific days of euthanasia and necropsy endpoints (*) are indicated for each of the animals. No treatment was required for the kra monkeys.

**Supplemental Fig. 4.** Flow Cytometry Gating Strategy. Representative flow cytometry gating strategy for measuring erythroid lineage cells in bone marrow aspirate collected from rhesus and kra monkeys.

**Supplemental Fig. 5.** Characterization of the dynamics of RBC removal and production processes in a representative kra monkey (12C44) during a *P. knowlesi* infection. To quantify the haemodynamic processes during a *P. knowlesi* infection, a computational dynamical model was used that was previously developed to faithfully track the blood dynamics in *Plasmodium* infected monkeys [57–59]. The model was formulated as a set of discrete recursive equations, where the pools of reticulocytes, RBCs, and iRBCs were stratified into age classes. The model directly represents that reticulocytes are released from the bone marrow with a certain age and rate, circulate for a day and then mature into RBCs. Pertinent model results (lines) are superimposed on experimental data (symbols). Shown are the circulating reticulocytes (A), mature RBCs (C), and infected RBCs (F), from which the model allowed the quantification of different causes of RBC removal (D and E). The NHP RBCs normally die after about 100 days due to senescence, or on a daily basis due to “random” effects, such as shear stresses (E). During *Plasmodium* infections, some of the healthy RBCs are also infected by merozoites and destroyed when the parasites are released (parasitization, D) or lost to a bystander effect (D). Interestingly, large numbers of RBCs were lost during the infection due to the bystander mechanism (D). The profile of RBCs (C) demonstrates that the kra monkey had severe anaemia and responded appropriately by increasing the erythropoietic output and releasing younger reticulocytes, thereby increasing the reticulocyte maturation time in circulation (B).

## DECLARATIONS

### ETHICS APPROVAL AND CONSENT TO PARTICIPATE

All experimental, surgical, and necropsy procedures were approved by Emory’s IACUC and the Animal Care and Use Review Office (ACURO) of the US Department of Defense and followed accordingly. The Emory’s IACUC approval number was PROTO201700484 - YER-2003344- ENTRPR-A.

### CONSENT FOR PUBLICATION

Not applicable

### AVAILBILITY OF DATA AND MATERIALS

All clinical and telemetry data were rigorously validated and quality controlled. The majority of data analyzed here, protocols and extensive metadata are publicly available in PlasmoDB [61]: http://plasmodb.org/plasmo/mahpic.jsp

http://plasmodb.org/common/downloads/MaHPIC/Experiment_07/

http://plasmodb.org/common/downloads/MaHPIC/Experiment_33/

http://plasmodb.org/common/downloads/MaHPIC/Experiment_35/

Data not available through public data repositories are included as supplementary files (e.g., tissue parasite counts are included in S1 Spreadsheet). Data not in public databases or supplementary files (e.g., flow cytometry data files) can be requested directly from the corresponding author.

## AUTHOR CONTRIBUTIONS

Conceived and designed the experiments: MSP, CJJ, JAB, JSW, LLF, RT, JCK, AM, SG, EOV, JBG, RJC, MRG, and members of the MaHPIC-Consortium. Performed the experiments: MSP, CJJ, JAB, JSW, MCM, CLS, LLF, WTC, JJ, SAL, SRS, AH, DM, EK. Performed data analysis: MSP, CJJ, JAB, LLF, SG, JBG. Interpreted the data analysis: MSP, CJJ, RJC, MRG and members of the MaHPIC-Consortium. Managed and led validation and quality control of datasets for clinical and telemetry results and deposited the data and metadata: MVN, JH, JDD, JCK. Generated the figures: MSP, CJJ, LLF, SG. Wrote the paper: MSP, CCJ, MRG. Provided manuscript editorial contributions: LLF, AM, SG, EOV, RJC. All authors reviewed and approved submission of this manuscript.

## ACKNOWLEDGEMENTS

The authors thank John W. Barnwell for discussions, bringing knowledge of *P. knowlesi* infections and malaria to this project and the provision of cryopreserved sporozoites. Elizabeth Strobert is thanked for consultations and advice on animal protocols. E-van Dessasau is thanked for technical assistance in preparing and staining histopathological slides. The YNPRC staff are acknowledged for assistance with procedures involving NHPs, and the Emory Pediatric/Winship Flow Cytometry and Yerkes Flow Cytometry Cores are recognized for maintaining core facilities made available in this project. *MaHPIC Consortium Members*. MaHPIC members participating in discussions at the time of the planning, implementation, or analysis of this project include: Dave C. Anderson, Ferhat Ay, Cristiana F. A. Brito, John W. Barnwell, Megan DeBarry, Steven E. Bosinger, Jung-Ting Chien, Jinho Choi, Anuj Gupta, Chris Ibegbu, Xuntian Jiang, Dean P. Jones, Nicolas Lackman, Tracey J. Lamb, Frances E.-H. Lee, Karine Gaelle Le Roche, Shuzhao Li, Esmeralda V.S. Meyer, Diego M. Moncada-Giraldo, Dan Ory, Jan Pohl, Saeid Safaei, Igñacio Sanz, Maren Smith, Gregory Tharp, ViLinh Tran, Elizabeth D. Trippe, Karan Uppal, Susanne Warrenfeltz, Tyrone Williams, Zerotti L. Woods.

## AUTHOR’S INFORMATION

Not Applicable

