## Supplementary figures and images for "Clinical recovery of *Macaca fascicularis* infected with *Plasmodium knowlesi*"

### Supplemental Fig 1

#### E07: Longitudinal *P. knowlesi* Infection in Kra Monkeys

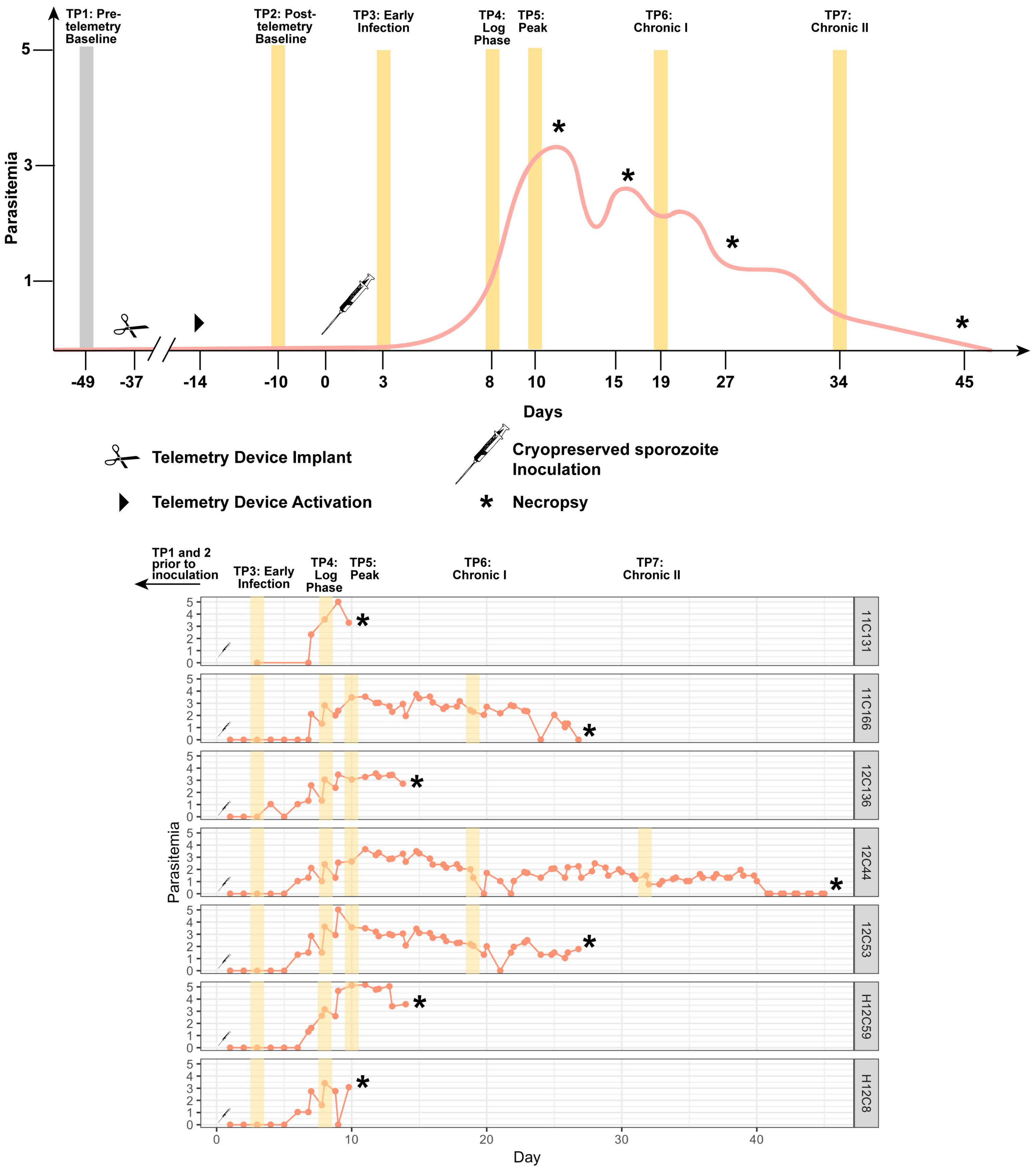

### Supplemental Fig 2

#### Acute *P. knowlesi* Infection in Kra

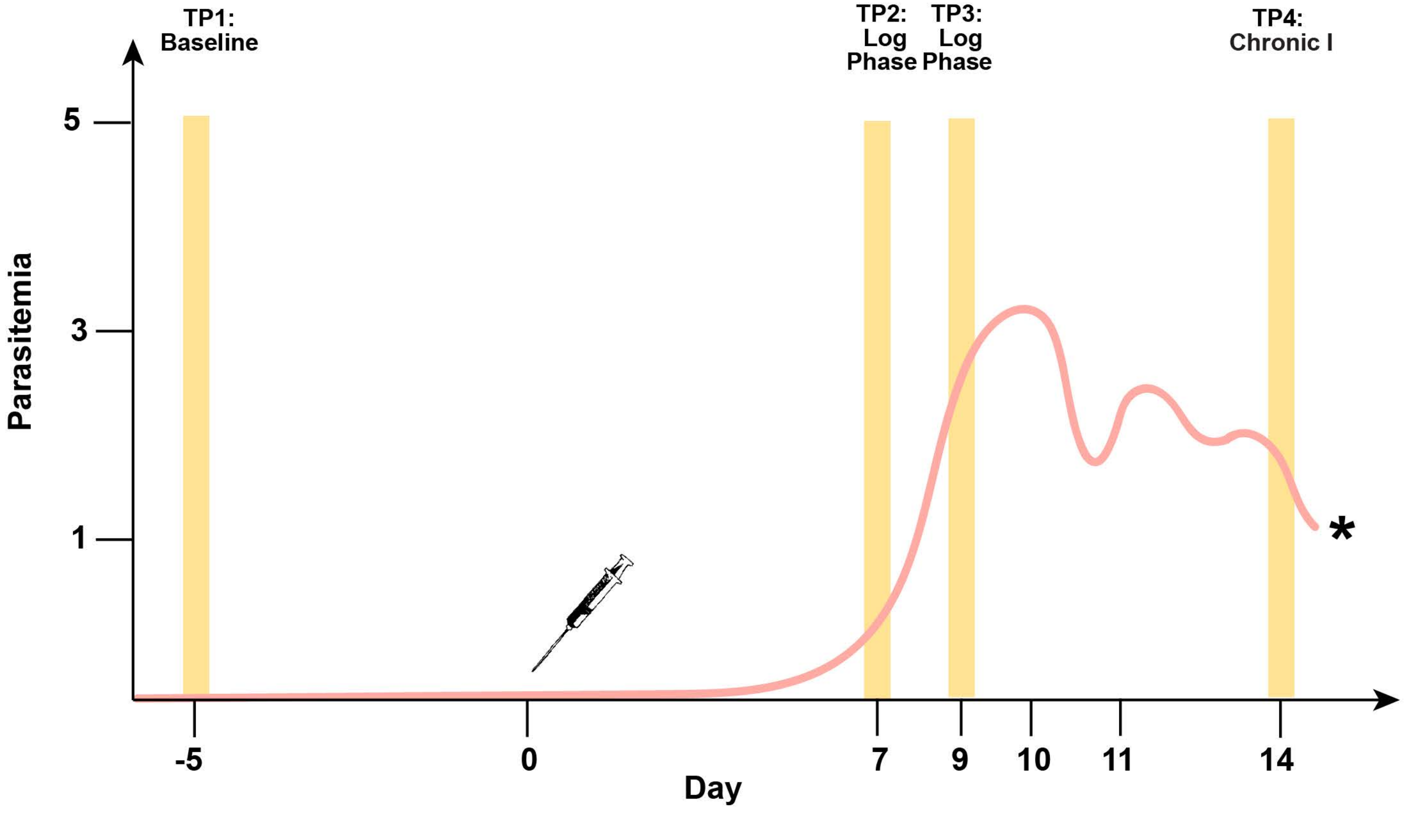


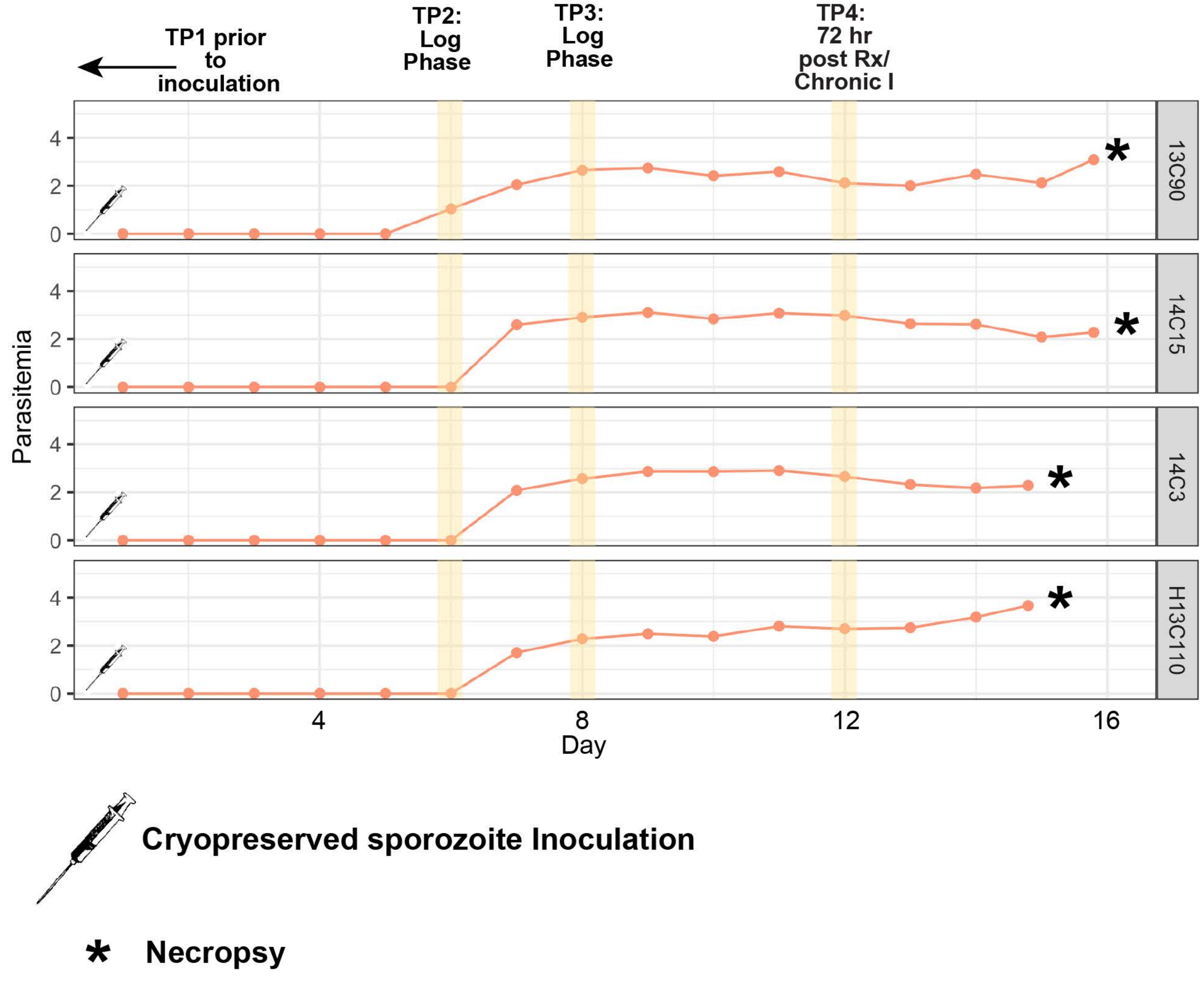

### Supplemental Fig 3

#### E35: Chronic *P. knowlesi* Infection in Kra

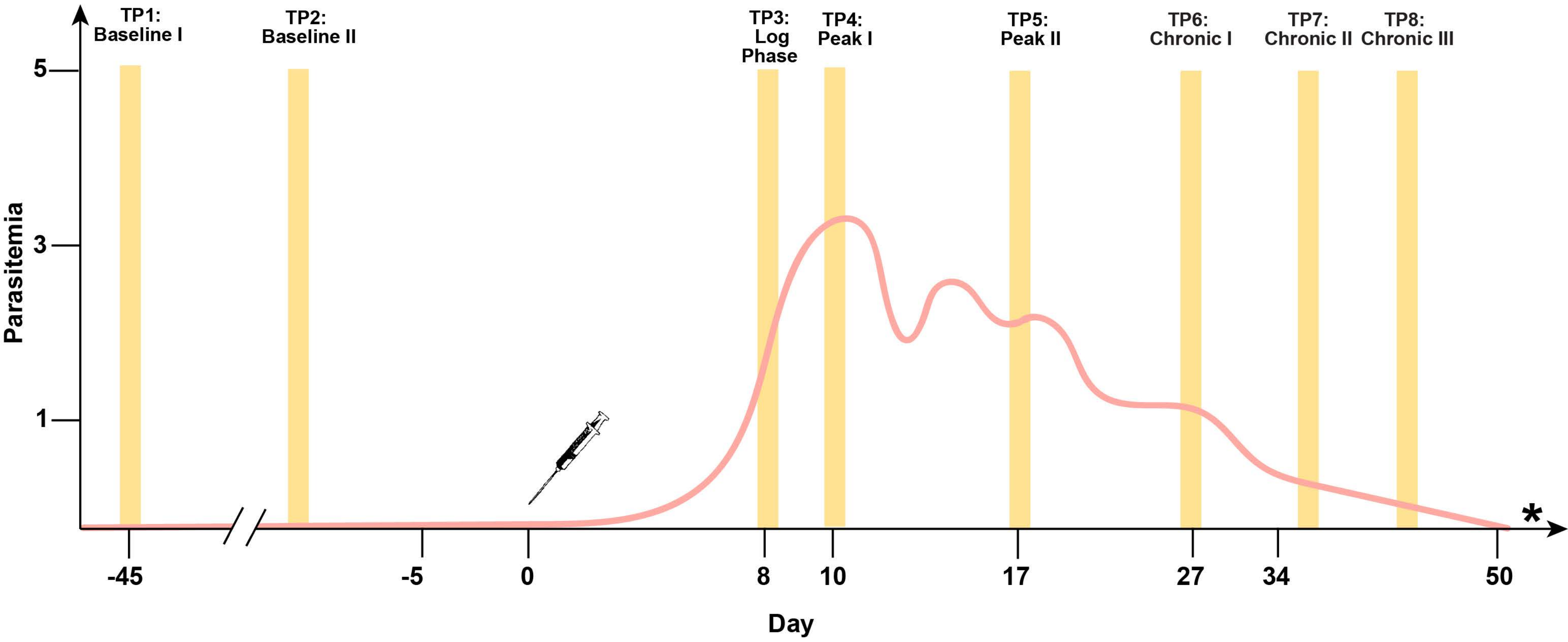


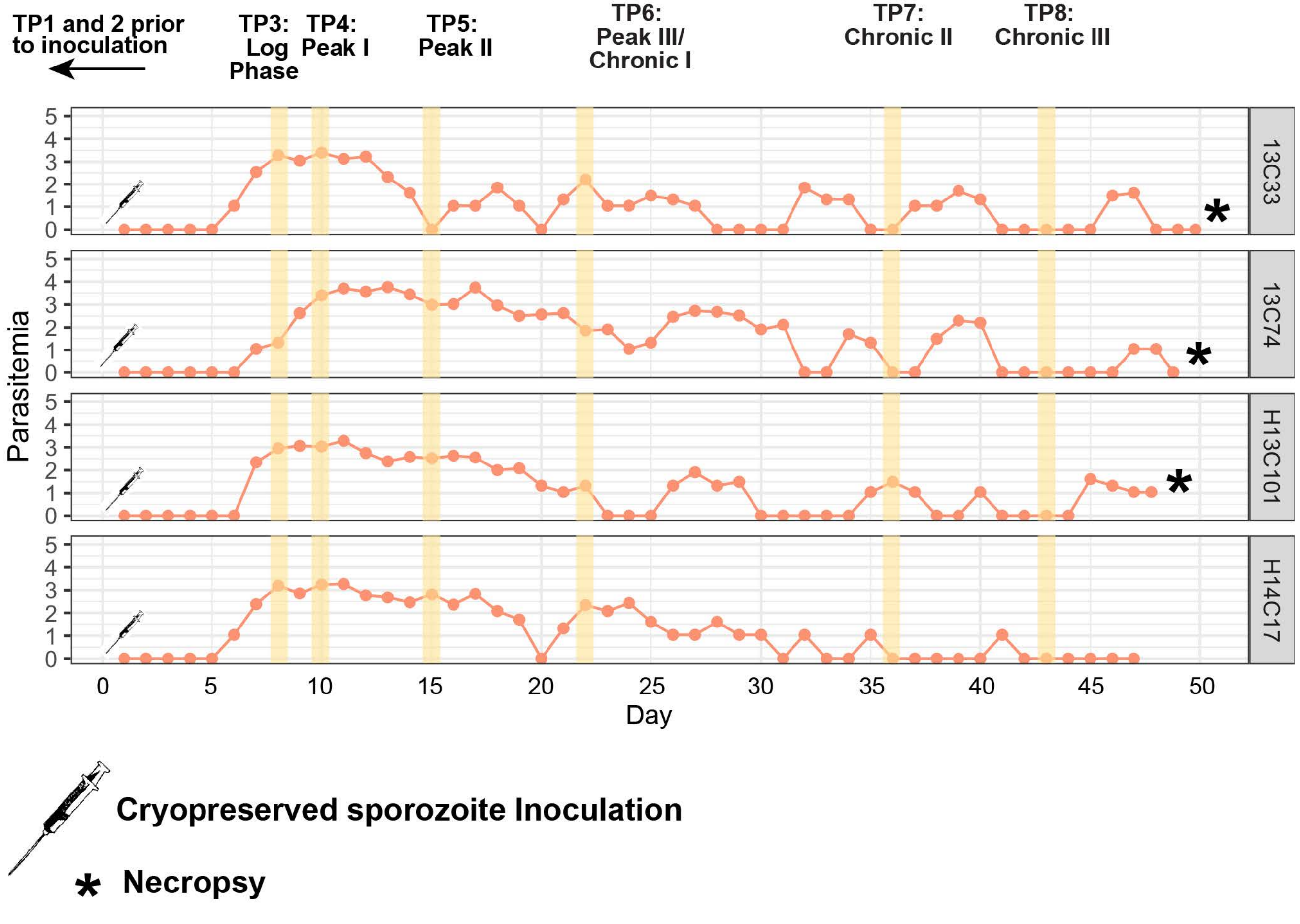

### Supplemental Fig 4

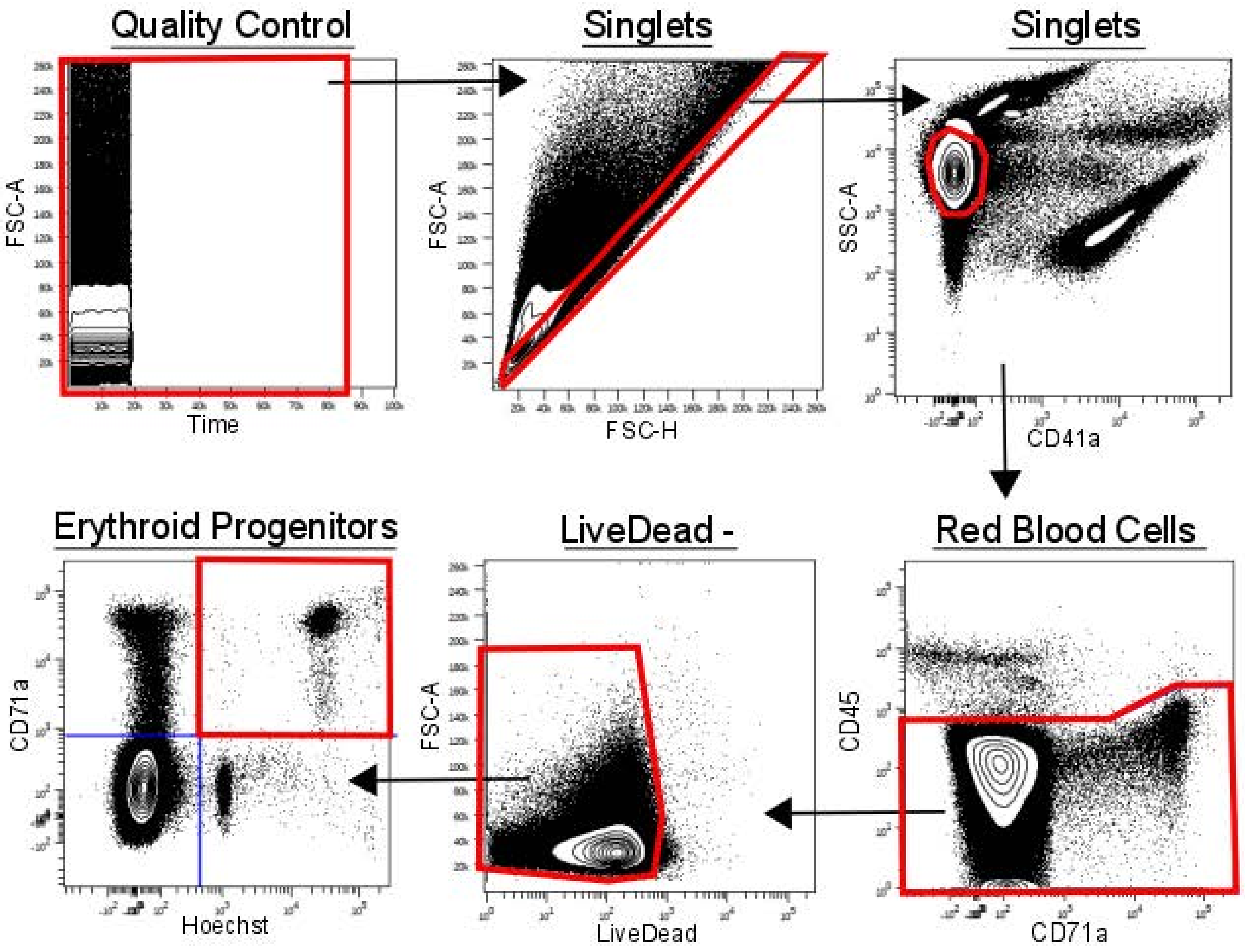

### Supplemental Fig 5

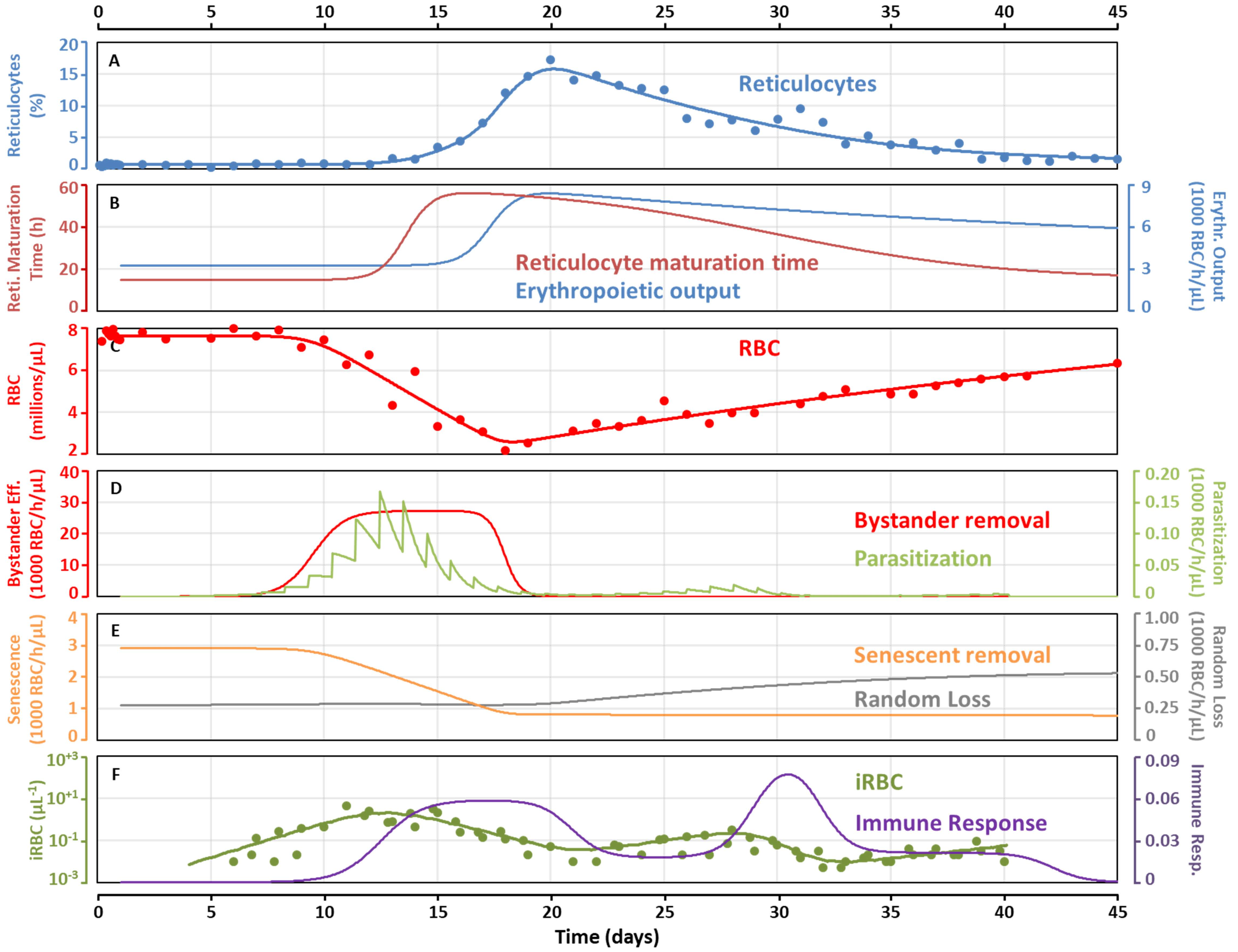
