## Supplemental Table 1 for "Clinical recovery of *Macaca fascicularis* infected with *Plasmodium knowlesi*"

| **Supplemental Table 1: *Macaca fascicularis* Cohorts** | | | | |
| --- | --- | --- | --- | --- |
| **Cohort Number** (Experiment Number)^§^ | **Infection Type** | **Monkey Codes** | **Brief Experimental Description**  (*Supplementary Figs. 1-3 describe and show graphed summaries*) | **Telemetry**  **Implants** |
| **1** (E07) | Pilot Experiment: *P. knowlesi*  Acute & Chronic  Infection of  *M. fascicularis* | 11C131, 11C166, 12C36, 12C44, 12C53, H12C59, H12C8 | Seven *M. fascicularis* were inoculated with *P. knowlesi* sporozoites obtained from fresh mosquito salivary gland dissections, but for unexplained reasons parasitaemias did not develop. The monkeys were subsequently infected with cryopreserved *P. knowlesi* sporozoites (the same batch used for all other experiments in this table). Blood and bone marrow samples were collected for analysis throughout the course of the infections and euthanasia and necropsies for pathology analyses were performed at pre-determined times. | Yes  *Continuous telemetry data were collected*^†^ |
| **2** (E33) | Iterative Experiment:  *P. knowlesi*  Acute Infections & Establishment of Chronic Infections | 13C90, 14C15, 14C3, H13C110, | Four *M. fascicularis* were infected with cryopreserved *P. knowlesi* sporozoites. Blood and bone marrow samples were collected for analysis at pre-determined intervals. The animals were euthanised and necropsied for pathology analyses within 2-3 weeks of inoculation, as the *M. fascicularis* were controlling their infections. | No |
| **3** (E34) | Control  Data | 13C102,13C105,  13C129 | Three *M. fascicularis* were sacrificed to provide normal control samples for analysis in conjunction with samples from infected macaques. | No |
| **4** (E35) | Iterative Experiment:  *P. knowlesi*  Acute & Chronic  Infection | 13C33, 13C74, H13C101, H14C17 | Four *M. fascicularis* were infected with cryopreserved *P. knowlesi* sporozoites. Blood and bone marrow samples were collected for analysis at predetermined intervals throughout the experiment. The animals were euthanised and necropsied for pathology analyses between days 48-50. | No |

**S1 Table.** **Macaque Cohort and Experimental Summaries.** Details regarding four monkey cohorts involved in the current study are summarised, three of which were experimentally infected with *P. knowlesi* sporozoites, while one served as control group. The cohorts are listed in the order that experiments using these animals were performed. These animals and their longitudinal infection designs were part of a systems biology program, with iterative cohort experimentation designed to satisfy the goals of those research programs. ^§^The non-sequential experimental numbering (E07, E33, E34, and E35) reflects the experimental numbers assigned in the MaHPIC Laboratory Information Management System. ^†^Telemetry data were collected for temperature, blood pressure, heart rate and activity level (manuscript in preparation).
