## Supplemental Table 2 for "Clinical recovery of *Macaca fascicularis* infected with *Plasmodium knowlesi*"

| **Supplemental Table 2 Summary of histopathology scores for *P. knowlesi*-infected macaques** | | | | | | | | | | | | | | | |
| --- | --- | --- | --- | --- | --- | --- | --- | --- | --- | --- | --- | --- | --- | --- | --- |
|  | **Monkey** | | | | | | | | | | | | | | |
| **Histopathology** | 11C131 | 11C166 | 12C44 | 12C53 | 12C136 | H12C8 | H12C59 | 13C90 | 13C33 | 14C3 | 14C15 | H13C101 | H13C110 | H14C17 | 13C74 |
| **Colon** |  |  |  |  |  |  |  |  |  |  |  |  |  |  |  |
| *Inflammation* | 2 | 0 | 1 | 1 | 1 | 1 | 1 | 1 | 1 | 1 | 1 | 1 | 1 | 1 | 1 |
| *Oedema* | 0 | 0 | 0 | 0 | 0 | 1 | 0 | 0 | 0 | 0 | 0 | 0 | 0 | 0 | 0 |
| **Duodenum** |  |  |  |  |  |  |  |  |  |  |  |  |  |  |  |
| *Inflammation* | 2 | 0 | 2 | 1 | 1 | 1 | 2 | 2 | 2 | 2 | 2 | 2 | 2 | 2 | 2 |
| **Jejunum** |  |  |  |  |  |  |  |  |  |  |  |  |  |  |  |
| *Inflammation* | 1 | 1 | 1 | 1 | 1 | 1 | 1 | 2 | 1 | 1 | 1 | 1 | 1 | 1 | 1 |
| **Stomach** |  |  |  |  |  |  |  |  |  |  |  |  |  |  |  |
| *Inflammation* | 2 | 2 | 2 | 2 | 0 | 1 | 0 | 1 | 3 | 2 | 0 | 4 | 2 | 0 | 0 |
| *Crypt Inflammation* | 0 | 0 | 0 | 0 | 0 | 0 | 0 | 0 | 0 | 0 | 0 | 0 | 0 | 0 | 0 |
| **Kidney** |  |  |  |  |  |  |  |  |  |  |  |  |  |  |  |
| *Inflammation* | 0 | 1 | 0 | 0 | 0 | 0 | 0 | 0 | 0 | 0 | 0 | 0 | 0 | 0 | 0 |
| *Haemorrhage* | 0 | 0 | 1 | 0 | 0 | 0 | 0 | 0 | 0 | 1 | 0 | 0 | 1 | 0 | 0 |
| *Tubular Degeneration* | 0 | 0 | 0 | 0 | 0 | 0 | 2 | 0 | 0 | 0 | 0 | 0 | 2 | 1 | 0 |
| *Glomerular Hypercellularity* | 1 | 1 | 1 | 1 | 1 | 1 | 1 | 1 | 1 | 1 | 1 | 1 | 1 | 1 | 1 |
| **Liver** |  |  |  |  |  |  |  |  |  |  |  |  |  |  |  |
| *Inflammation* | 2 | 2 | 2 | 2 | 2 | 2 | 2 | 2 | 2 | 2 | 2 | 1 | 2 | 2 | 1 |
| *Kupffer Cell Hyperplasia* | 2 | 2 | 2 | 2 | 3 | 3 | 2 | 2 | 2 | 2 | 2 | 1 | 2 | 2 | 2 |
| **Lung** |  |  |  |  |  |  |  |  |  |  |  |  |  |  |  |
| *Inflammation* | 0 | 0 | 0 | 0 | 0 | 0 | 0 | 0 | 0 | 0 | 0 | 0 | 0 | 1 | 2 |
| *Haemorrhage* | 0 | 0 | 0 | 0 | 0 | 0 | 0 | 1 | 2 | 0 | 0 | 1 | 0 | 0 | 1 |
| *Hyperplasia* | 1 | 0 | 1 | 1 | 0 | 2 | 1 | 1 | 1 | 1 | 1 | 1 | 1 | 1 | 2 |
| *Fibrosis* | 2 | 1 | 2 | 1 | 0 | 2 | 2 | 2 | 2 | 2 | 2 | 3 | 2 | 3 | 2 |
| **Ventricle** |  |  |  |  |  |  |  |  |  |  |  |  |  |  |  |
| *Inflammation* | 2 | 0 | 0 | 1 | 1 | 0 | 0 | 0 | 0 | 0 | 0 | 0 | 0 | 0 | 0 |
| *Oedema* | 2 | 0 | 0 | 1 | 0 | 0 | 0 | 0 | 0 | 0 | 0 | 0 | 0 | 0 | 0 |
| *Haemorrhage* | 0 | 0 | 0 | 0 | 2 | 0 | 0 | 0 | 0 | 0 | 0 | 0 | 0 | 0 | 0 |

**Supplemental Table 2 Summary of histopathology scores for *P. knowlesi*-infected macaques.** Nineteen tissues were examined by a veterinary pathologist and scored semi-quantitatively in the categories above. Spleen, lymph nodes, and bone marrow were not scored because the changes were deemed to be physiological reactions to infection and/or anaemia. The following tissues were examined for histopathological changes but omitted from this table because no changes were found: aorta, adrenal gland, cerebrum, cerebellum, eye, midbrain, muscle, skin, testis, thymus, and omentum. These tissues were assigned a score of zero for modeling purposes.
