## Supplemental Table 3 for "Clinical recovery of *Macaca fascicularis* infected with *Plasmodium knowlesi*"

**Supplemental Table 3: Linear Regression: Spleen Weight *vs*. Time of Infection**

|  | Estimate | Std. Error | *t* value | *p*-value | Significance |
| --- | --- | --- | --- | --- | --- |
| Intercept | 0.572 | 0.080 | 7.145 | 7.54x10^-6^ | *** |
| Necropsy Day | -0.004 | 0.003 | -1.430 | 0.176 | NS |
| *n* = 15; *df* = 13; Adjusted *R*^2^ = 0.069; *F*-statistic = 2.045; *p* = 0.176 | | | | | |
