## Supplemental Table 4 for "Clinical recovery of *Macaca fascicularis* infected with *Plasmodium knowlesi*"

| **Supplemental Table 4: Tukey HSD Post-Hoc Pairwise Comparison of Histopathological Tissue Score** | | | | | | |
| --- | --- | --- | --- | --- | --- | --- |
| Organ 1 (I) | Organ 2 (J) | Mean Difference (I-J) | Adjusted P-value | Significance | LB | UB |
| Adrenal Gland | Aorta | -1.465x10-14 | 1.000 | NS | -1.247 | 1.247 |
|  | Cerebellum | -1.643x10-14 | 1.000 | NS | -1.247 | 1.247 |
|  | Cerebrum | -1.488x10-14 | 1.000 | NS | -1.247 | 1.247 |
|  | Colon | 1.000 | 0.303 | NS | -0.247 | 2.266 |
|  | Duodenum | 1.500 | 0.004 | ** | 0.253 | 2.747 |
|  | Eye | 1.577x10-14 | 1.000 | NS | -1.247 | 1.247 |
|  | Jejunum | 1.100 | 0.160 | NS | -0.147 | 2.347 |
|  | Kidney | 1.700 | < 0.005 | *** | 0.453 | 2.947 |
|  | Liver | 3.900 | 0.000 | **** | 2.653 | 5.147 |
|  | Lung | 3.400 | 0.000 | **** | 2.153 | 4.647 |
|  | Midbrain | -1.488x10-14 | 1.000 | NS | -1.247 | 1.247 |
|  | Muscle | -1.632x10-14 | 1.000 | NS | -1.247 | 1.247 |
|  | Omentum | 1.565x10-14 | 1.000 | NS | -1.247 | 1.247 |
|  | Skin | -1.554x10-14 | 1.000 | NS | -1.247 | 1.247 |
|  | Stomach | 1.600 | 0.001 | ** | 0.353 | 2.847 |
|  | Testis | -1.432x10-14 | 1.000 | NS | -1.247 | 1.247 |
|  | Thymus | -1.532x10-14 | 1.000 | NS | -1.247 | 1.247 |
|  | Ventricle | 0.900 | 0.501 | NS | -0.347 | 2.147 |
| Aorta | Cerebellum | -1.776x10-15 | 1.000 | NS | -1.247 | 1.247 |
|  | Cerebrum | -2.220x10-16 | 1.000 | NS | -1.247 | 1.247 |
|  | Colon | 1.000 | 0.303 | NS | -0.247 | 2.247 |
|  | Duodenum | 1.500 | 0.004 | ** | 0.253 | 2.747 |
|  | Eye | -1.110x10-15 | 1.000 | NS | -1.247 | 1.247 |
|  | Jejunum | 1.100 | 0.160 | NS | -0.147 | 2.347 |
|  | Kidney | 1.700 | <0.005 | *** | 0.453 | 2.947 |
|  | Liver | 3.900 | 0.000 | **** | 2.653 | 5.147 |
|  | Lung | 3.400 | 0.000 | **** | 2.153 | 4.647 |
|  | Midbrain | -2.220x10-16 | 1.000 | NS | -1.247 | 1.247 |
|  | Muscle | -1.665x10-16 | 1.000 | NS | -1.247 | 1.247 |
|  | Omentum | -9.992x10-16 | 1.000 | NS | -1.247 | 1.247 |
|  | Skin | -8.882x10-16 | 0.0000 | NS | -1.247 | 1.247 |
|  | Stomach | 1.600 | 0.001 | ** | 0.353 | 2.847 |
|  | Thymus | 3.331x10-16 | 1.000 | NS | -1.247 | 1.247 |
|  | Ventricle | -6.661x10-16 | 1.000 | NS | -1.247 | 1.247 |
| Cerebellum | Cerebrum | 1.554x10-15 | 1.000 | NS | -1.247 | 1.247 |
|  | Colon | 1.000 | .303 | NS | -0.247 | 2.247 |
|  | Duodenum | 1.500 | 0.004 | ** | 0.253 | 2.747 |
|  | Eye | 6.661x10-16 | 1.000 | NS | -1.247 | 1.247 |
|  | Jejunum | 1.100 | 0.160 | NS | -0.147 | 2.347 |
|  | Kidney | 1.700 | <0.005 | *** | 0.453 | 2.947 |
|  | Liver | 3.900 | 0.000 | **** | 2.653 | 5.147 |
|  | Lung | 2.400 | 0.000 | **** | 2.153 | 4.647 |
|  | Midbrain | 1.554x10-15 | 1.000 | NS | -1.247 | 1.247 |
|  | Muscle | 1.110x10-16 | 1.000 | NS | -1.247 | 1.247 |
|  | Omentum | 7.772x10-16 | 1.000 | NS | -1.247 | 1.247 |
|  | Skin | 8.882x10-16 | 1.000 | NS | -1.247 | 1.247 |
|  | Stomach | 1.600 | 0.001 | ** | 0.353 | 2.847 |
|  | Testis | 2.109x10-15 | 1.000 | NS | -1.247 | 1.247 |
|  | Thymus | 1.110x10-15 | 1.000 | NS | -1.247 | 1.247 |
|  | Ventricle | 0.900 | 0.501 | NS | -0.347 | 2.147 |
| Cerebrum | Colon | 1.00 | 0.303 | NS | -0.247 | 2.247 |
|  | Duodenum | 1.500 | 0.004 | ** | -0.253 | 2.747 |
|  | Eye | -8.882x10-16 | 1.0000 | NS | -1.247 | 1.247 |
|  | Jejunum | 1.100 | 0.000 | **** | -0.147 | 2.347 |
|  | Kidney | 1.700 | <0.005 | ** | 0.453 | 2.947 |
|  | Liver | 3.900 | 0.000 | **** | 2.653 | 5.147 |
|  | Lung | 3.400 | 0.000 | **** | 2.153 | 4.647 |
|  | Midbrain | 1.554x10-15 | 1.0000 | NS | -1.247 | 1.247 |
|  | Muscle | 1,1102x10-16 | 1.0000 | NS | -1.247 | 1.247 |
|  | Omentum | 7.772x10-16 | 1.0000 | NS | -1.247 | 1.247 |
|  | Skin | 8.882x10-16 | 1.0000 | NS | -1.247 | 1.247 |
|  | Stomach | 1.600 | 0.001 | ** | 0.353 | 2.847 |
|  | Testis | 2.109x10-15 | 1.0000 | NS | -1.247 | 1.247 |
|  | Thymus | 1.110x10-15 | 1.0000 | NS | -1.247 | 1.247 |
|  | Ventricle | 0.900 | 0.501 | NS | -0.347 | 2.147 |
| Colon | Duodenum | 0.500 | 0.995 | NS | -0.747 | 1.747 |
|  | Eye | -1.000 | 0.303 | NS | -2.247 | 0.247 |
|  | Jejunum | 0.100 | 1.000 | NS | -1.147 | 1.347 |
|  | Kidney | 0.700 | 0.880 | NS | -0.547 | 1.947 |
|  | Liver | 2.900 | 0.000 | **** | 1.653 | 4.147 |
|  | Lung | 2.400 | 0.000 | **** | 1.153 | 3.647 |
|  | Midbrain | -1.000 | 0.303 | NS | -2.247 | 0.247 |
|  | Muscle | -1.000 | 0.303 | NS | -2.247 | 0.247 |
|  | Omentum | -1.000 | 0.303 | NS | -2.247 | 0.247 |
|  | Skin | -1.000 | 0.303 | NS | -2.247 | 0.247 |
|  | Stomach | 0.600 | 0.968 | NS | -0.647 | 1.847 |
|  | Testis | -1.000 | 0.303 | NS | -2.247 | 0.247 |
|  | Thymus | -1.000 | 0.303 | NS | -2.247 | 0.247 |
|  | Ventricle | -0.100 | 1.000 | NS | -1.347 | 1.147 |
| Duodenum | Eye | -1.500 | 0.004 | ** | -2.747 | -0.253 |
|  | Jejunum | -0.400 | 1.000 | NS | -1.647 | 0.847 |
|  | Kidney | 0.200 | 1.000 | NS | -1.047 | 1.447 |
|  | Liver | 2.400 | 0.000 | **** | 1.153 | 3.647 |
|  | Lung | 1.900 | <0.005 | *** | 0.653 | 3.147 |
|  | Midbrain | -1.500 | 0.004 | ** | -2.747 | -0.253 |
|  | Muscle | -1.500 | 0.004 | ** | -2.747 | -0.253 |
|  | Omentum | -1.500 | 0.004 | ** | -2.747 | -0.253 |
|  | Skin | -1.500 | 0.004 | ** | -2.747 | -0.253 |
|  | Stomach | 0.100 | 1.000 | NS | -1.147 | 1.347 |
|  | Testis | -1.500 | 0.004 | ** | -2.747 | -0.253 |
|  | Thymus | -1.500 | 0.004 | ** | -2.747 | -0.253 |
|  | Ventricle | -.600 | 0.968 | NS | -1.847 | 0.647 |
| Eye | Jejunum | 1.100 | 0.160 | NS | -0.147 | 2.347 |
|  | Kidney | 1.700 | < 0.005 | *** | 0.453 | 2.947 |
|  | Liver | 3.900 | 0.000 | **** | 2.653 | 5.147 |
|  | Lung | 3.400 | 0.000 | **** | 2.153 | 4.647 |
|  | Midbrain | 8.882x10-16 | 1.000 | NS | -1.247 | 1.247 |
|  | Muscle | -5.551x10-16 | 1.000 | NS | -1.247 | 1.247 |
|  | Omentum | 1.110x10-16 | 1.000 | NS | -1.247 | 1.247 |
|  | Skin | 2.220x10-16 | 1.000 | NS | -1.247 | 1.247 |
|  | Stomach | 1.600 | 0.001 | ** | 0.353 | 2.847 |
|  | Testis | 1.443x10-15 | 1.000 | NS | -1.247 | 1.247 |
|  | Thymus | 4.441x10-16 | 1.000 | NS | -1.247 | 1.247 |
|  | Ventricle | 0.900 | 0.501 | NS | -0.347 | 2.147 |
| Jejunum | Kidney | -0.600 | 0.968 | NS | -0.647 | 1.847 |
|  | Liver | 2.800 | 0.000 | **** | 1.553 | 4.047 |
|  | Lung | 2.300 | 0.000 | **** | 1.053 | 3.547 |
|  | Midbrain | -1.100 | 0.160 | NS | -2.347 | 0.147 |
|  | Muscle | -1.100 | 0.160 | NS | -2.347 | 0.147 |
|  | Omentum | -1.100 | 0.160 | NS | -2.347 | 0.147 |
|  | Skin | -1.100 | 0.160 | NS | -2.347 | 0.147 |
|  | Stomach | 0.500 | 0.995 | NS | -0.747 | 1.747 |
|  | Testis | -1.100 | 0.160 | NS | -2.347 | 0.147 |
|  | Thymus | -1.100 | 0.160 | NS | -2.347 | 0.147 |
|  | Ventricle | -0.200 | 1.000 | NS | -1.447 | 1.047 |
| Kidney | Liver | 2.200 | 0.000 | **** | 0.953 | 3.447 |
|  | Lung | 1.700 | < 0.005 | *** | 0.453 | 2.947 |
|  | Midbrain | -1.700 | < 0.005 | *** | -2.947 | -0.453 |
|  | Muscle | -1.700 | < 0.005 | *** | -2.947 | -0.453 |
|  | Omentum | -1.700 | < 0.005 | *** | -2.947 | -0.453 |
|  | Skin | -1.700 | < 0.005 | *** | -2.947 | -0.453 |
|  | Stomach | -0.100 | 1.000 | NS | -1.347 | 1.147 |
|  | Testis | -1.700 | < 0.005 | *** | -2.947 | -0.453 |
|  | Thymus | -1.700 | < 0.005 | *** | -2.947 | -0.453 |
|  | Ventricle | -0.800 | 0.713 | NS | -2.047 | 0.447 |
| Liver | Lung | -0.500 | 0.995 | NS | -1.747 | 0.747 |
|  | Midbrain | -3.900 | 0.000 | **** | -5.147 | -2.653 |
|  | Muscle | -3.900 | 0.000 | **** | -5.147 | -2.653 |
|  | Omentum | -3.900 | 0.000 | **** | -5.147 | -2.653 |
|  | Skin | -3.900 | 0.000 | **** | -5.147 | -2.653 |
|  | Stomach | -2.300 | 0.000 | **** | -3.547 | -1.053 |
|  | Testis | -3.900 | 0.000 | **** | -5.147 | -2.653 |
|  | Thymus | -3.900 | 0.000 | **** | -5.147 | -2.653 |
|  | Ventricle | -3.000 | 0.000 | **** | -4.247 | -1.753 |
| Lung | Midbrain | -3.400 | 0.000 | **** | -4.647 | -2.153 |
|  | Muscle | -3.400 | 0.000 | **** | -4.647 | -2.153 |
|  | Omentum | -3.400 | 0.000 | **** | -4.647 | -2.153 |
|  | Skin | -3.400 | 0.000 | **** | -4.647 | -2.153 |
|  | Stomach | -1.800 | < 0.005 | *** | -3.047 | -0.553 |
|  | Testis | -3.400 | 0.000 | **** | -4.647 | -2.153 |
|  | Thymus | -3.400 | 0.000 | **** | -4.647 | -2.153 |
|  | Ventricle | -2.500 | 0.000 | **** | -3.747 | -1.253 |
| Midbrain | Muscle | -1.443x10-15 | 1.000 | NS | -1.247 | 1.247 |
|  | Omentum | -7.772x10-16 | 1.000 | NS | -1.247 | 1.247 |
|  | Skin | -6.661x10-16 | 1.000 | NS | -1.247 | 1.247 |
|  | Stomach | 1.600 | 0.001 | ** | 0.353 | 2.847 |
|  | Testis | 5.552x10-16 | 1.000 | NS | -1.247 | 1.247 |
|  | Thymus | -4.441x10-16 | 1.000 | NS | -1.247 | 1.247 |
|  | Ventricle | 0.900 | 0.501 | NS | -0.347 | 2.147 |
| Muscle | Omentum | 6.661x10-16 | 1.000 | NS | -1.247 | 1.247 |
|  | Skin | 7.772x10-16 | 1.000 | NS | -1.247 | 1.247 |
|  | Stomach | 1.600 | 0.001 | ** | 0.353 | 2.847 |
|  | Testis | 1.998x10-15 | 1.000 | NS | -1.247 | 1.247 |
|  | Thymus | 9.992x10-16 | 1.000 | NS | -1.247 | 1.247 |
|  | Ventricle | 0.900 | 0.501 | NS | -0.347 | 2.147 |
| Omentum | Skin | 1.110x10-16 | 1.000 | NS | -1.247 | 1.247 |
|  | Stomach | 1.600 | 0.001 | ** | 0.353 | 2.847 |
|  | Testis | 1.332x10-15 | 1.000 | NS | -1.247 | 1.247 |
|  | Thymus | 3.330x10-16 | 1.000 | NS | -1.247 | 1.247 |
|  | Ventricle | 0.900 | 0.501 | NS | -0.347 | 2.147 |
| Skin | Stomach | 1.600 | 0.001 | ** | 0.353 | 2.847 |
|  | Testis | 1.221x10-15 | 1.000 | NS | -1.247 | 1.247 |
|  | Thymus | 2.220x10-16 | 1.000 | NS | -1.247 | 1.247 |
|  | Ventricle | 0.900 | 0.501 | NS | -0.347 | 2.147 |
| Stomach | Testis | -1.600 | 0.001 | ** | -2.847 | -0.353 |
|  | Thymus | -1.600 | 0.001 | ** | -2.847 | -0.353 |
|  | Ventricle | -0.700 | 0.880 | NS | -1.947 | 0.547 |
| Testis | Thymus | -9.992x10-16 | 1.000 | NS | -1.247 | 1.247 |
|  | Ventricle | 0.900 | 0.501 | NS | -0.347 | 2.147 |
| Ventricle | Thymus | 0.900 | 0.501 | NS | -0.347 | 2.147 |
