## Supplemental Table 5 for "Clinical recovery of *Macaca fascicularis* infected with *Plasmodium knowlesi*"

**Supplemental Table 5: Linear Regression: Pathology Score *vs*. Necropsy Day**

|  | | Estimate | Std. Error | *t* value | *p*-value | Significance |
| --- | --- | --- | --- | --- | --- | --- |
| Intercept | | 0.693 | 0.208 | 3.332 | 0.001 | ** |
| Necropsy Day | | 0.003 | 0.006 | 0.560 | 0.576 | NS |
|  | *n*= 188; *df* = 190; Adjusted *R*^2^ = -0.004; *F*-statistic = 0.314; *p* = 0.576 | | | | | |
