## Supplemental Table 6 for "Clinical recovery of *Macaca fascicularis* infected with *Plasmodium knowlesi*"

**Supplemental Table 6. Summary Statistics for Parasite Tissue Counts**

| Tissue | Mean | Standard Deviation | Standard Error | *N* |
| --- | --- | --- | --- | --- |
| Adrenal Gland | 8.636 | 9.553 | 2.880 | 11 |
| Aorta | 1.818 | 3.545 | 1.069 | 11 |
| Bone Marrow | 11.368 | 27.941 | 6.410 | 19 |
| Cerebellum | 0.0833 | 0.289 | 0.083 | 12 |
| Cerebrum | 0.0833 | 0.289 | 0.083 | 12 |
| Colon | 9.909 | 20.186 | 6.086 | 11 |
| Duodenum | 6.636 | 8.958 | 2.701 | 11 |
| Eye | 2.727 | 6.828 | 2.059 | 11 |
| Jejunum | 3.273 | 4.692 | 1.415 | 11 |
| Kidney | 6.364 | 7.852 | 2.367 | 11 |
| Liver | 7.727 | 9.100 | 2.744 | 11 |
| Lung | 11.273 | 14.588 | 4.399 | 11 |
| Mesenteric LN | 1.364 | 2.248 | 0.678 | 11 |
| Midbrain | 0.000 | 0.000 | 0.000 | 11 |
| Muscle | 1.000 | 1.844 | 0.556 | 11 |
| Omentum | 1.091 | 1.514 | 0.456 | 11 |
| Skin | 0.583 | 0.900 | 0.260 | 12 |
| Spleen | 10.700 | 10.615 | 3.357 | 10 |
| Stomach | 6.818 | 9.358 | 2.821 | 11 |
| Testis | 0.727 | 1.489 | 0.449 | 11 |
| Thymus | 0.750 | 1.055 | 0.305 | 12 |
| Ventricle | 6.667 | 11.420 | 3.297 | 12 |
