## Supplemental Table 7 for "Clinical recovery of *Macaca fascicularis* infected with *Plasmodium knowlesi*"

| **Supplemental Table 7: Hierarchical Linear Regression Analysis** | | | | | | | | | | |
| --- | --- | --- | --- | --- | --- | --- | --- | --- | --- | --- |
|  | Model 1 | | | |  | Model 2 | | | |  |
|  | Estimate | Std. Error | t value | p-value | Significance | Estimate | Std. Error | t value | p-value | Significance |
| Intercept | 0.625 | 0.115 | 5.424 | 1.78x10^-7^ | *** | -0.013 | 0.255 | -0.051 | 0.959 | NS |
| Count | 0.030 | 0.011 | 2.817 | 0.005 | ** | -0.001 | 0.007 | 0.206 | 0.837 | NS |
| Organ |  |  |  |  |  |  |  |  |  |  |
| *Aorta* |  |  |  |  |  | 0.010 | 0.353 | 0.029 | 0.977 | NS |
| *Cerebellum* |  |  |  |  |  | 0.006 | 0.351 | 0.018 | 0.986 | NS |
| *Cerebrum* |  |  |  |  |  | 0.005 | 0.351 | 0.014 | 0.989 | NS |
| *Colon* |  |  |  |  |  | 0.998 | 0.350 | 2.851 | 0.005 | ** |
| *Duodenum* |  |  |  |  |  | 1.503 | 0.350 | 4.292 | 2.97x10^-5^ | *** |
| *Eye* |  |  |  |  |  | 0.002 | 0.350 | 0.005 | 0.996 | NS |
| *Jejunum* |  |  |  |  |  | 1.108 | 0.352 | 3.147 | 0.002 | ** |
| *Kidney* |  |  |  |  |  | 1.704 | 0.350 | 4.863 | 2.62x10^-6^ | **** |
| *Liver* |  |  |  |  |  | 3.901 | 0.350 | 11.147 | <2x10^-16^ | **** |
| *Lung* |  |  |  |  |  | 3.396 | 0.350 | 9.692 | <2x10^-16^ | **** |
| *Midbrain* |  |  |  |  |  | 0.004 | 0.350 | 0.011 | 0.992 | NS |
| *Muscle* |  |  |  |  |  | 0.012 | 0.354 | 0.032 | 0.974 | NS |
| *Omentum* |  |  |  |  |  | 0.011 | 0.355 | 0.032 | 0.974 | NS |
| *Skin* |  |  |  |  |  | 0.012 | 0.355 | 0.034 | 0.973 | NS |
| *Stomach* |  |  |  |  |  | 1.602 | 0.350 | 4.577 | 9.07x10^-6^ | **** |
| *Testis* |  |  |  |  |  | 0.012 | 0.355 | 0.034 | 0.973 | NS |
| *Thymus* |  |  |  |  |  | 0.901 | 0.350 | 2.574 | 0.011 | * |
| *Ventricle* |  |  |  |  |  | 0.001 | 0.007 | 0.206 | 0.837 | NS |
|  | N = 190; df = 188; Adjusted R^2^ = 0.035; F-statistic = 7.934;  p-value = 0.005 | | | | | N = 190; df = 170; Adjusted R^2^ = 0.678; F-statistic = 21.92;  p-value = <2.2x10^-16^ | | | | |
