## Supplemental Table 8 for "Clinical recovery of *Macaca fascicularis* infected with *Plasmodium knowlesi*"

| **Supplemental Table 8: Linear Regression** | | | | | | | | | | |  |  |  |  |  |  |  |  |  |  |
| --- | --- | --- | --- | --- | --- | --- | --- | --- | --- | --- | --- | --- | --- | --- | --- | --- | --- | --- | --- | --- |
|  | Model 1 | | | | | Model 2 | | | | | Model 3 | | | | | Model 4 | | | | |
|  | Est. | Std. Error | t value | p-value | Sig. | Est. | Std. Error | t value | p-value | Sig. | Est. | Std. Error | t value | p-value | Sig. | Est. | Std.  Error | t value | p-value | Sig. |
| Intercept | 0.625 | 0.115 | 5.424 | 1.78x10^-7^ | *** | 0.202 | 0.249 | 0.811 | 0.418 | NS | 0.250 | 0.350 | 0.714 | 0.476 | NS | 0.313 | 0.385 | 0.812 | 0.418 | NS |
| Count | 0.030 | 0.011 | 2.817 | 0.005 | ** | 0.039 | 0.012 | 3.370 | 9.12x10^-3^ | *** | 0.039 | 0.017 | 3.354 | 9.67x10^-4^ | *** | 0.040 | 0.012 | 3.369 | 9.19x10^-4^ | *** |
| DPI |  |  |  |  |  | 0.013 | 0.007 | 1.909 | 0.059 | NS | 0.116 | 0.009 | 1.366 | 0.173 | NS | 0.009 | 0.010 | 1.038 | 0.301 | NS |
| Parasitaemia |  |  |  |  |  |  |  |  |  |  | -2.322x10^-5^ | 1.190x10^-4^ | -0.195 | 0.846 | NS | 1.05x10^-4^ | 2.39x10^-4^ | -0.440 | 0.660 | NS |
| Cumulative Parasitaemia |  |  |  |  |  |  |  |  |  |  |  |  |  |  |  | 7.173x10^-7^ | 1.813x10^-6^ | 0.396 | 0.693 | NS |
|  | N = 190; df = 188; Adjusted R^2^ = 0.035; F-statistic = 7.934;  *p* = 0.005 | | | | | N = 190; df = 170; Adjusted R^2^ = 0.678; F-statistic = 21.92;  *p* = <2.2x10^-16^ | | | | | N = 190; df = 186; Adjusted R2 = 0.043; F-statistic = 3.889  *p* = 0.010 | | | | | N = 190; df = 185, Adjusted R2 = 0.040; F-statistic = 2.943  *p* = 0.022 | | | | |
