## Supplemental Table 9 for "Clinical recovery of *Macaca fascicularis* infected with *Plasmodium knowlesi*"

**S9 Table. Flow Antibody Cocktail**

| **Antibody** | **Fluorophore** | **Clone** | **Company** | **Titration (µl)** |
| --- | --- | --- | --- | --- |
| CD47 | FITC | MEM-122 | ThermoFisher Scientific | 10 |
| Band-3 | PE | BRIC-6 | ARP | 1 |
| CD41a | PE-Cy7 | HIP8 | Biolegend | 1.25 |
| CD71a | APC | LOI.1 | BD | 5 |
| Hoechst 33342 | Emission at 461 nm | N/A | BD | Final Concentration of 10 µg/ml |
| HLA-A,B,C | Alexfluor700 | W6/32 | Biolegend | 1.6 |
| LiveDead | Yellow | N/A | Life Technologies | 1 |
| CD45 | APC-Cy7 | D058-1283 | BD | 0.5 |
| Annexin V | BUV395 | N/A | BD | 5 |
